# Genome-wide parallelism in divergence and diversity across three Hydrophis sea snake species

**DOI:** 10.64898/2026.10.06.756852

**Authors:** Alastair J. Ludington, Kate L. Sanders

**Affiliations:** School of Biological sciences, Adelaide University, Adelaide, SA 5005, Australia; The South Australian Museum, Adelaide, Australia

**Keywords:** Sea snakes, genome scan, divergence landscape, adaptation, speciation, chemosensory

## Abstract

Understanding how genomic architecture and demographic history interact to shape patterns of genomic divergence is central to explaining adaptive radiation and speciation. Here, we provide the first full sequence patterns of diversity and differentiation for *Hydrophis* sea snakes, using population sampling of three ecomorphologically distinct, coexisting species that diverged near-simultaneously within the last few million years (*H. major, H. elegans* and *H. curtus*). We found recent population expansions in all three species, with little evidence for pervasive interspecific gene flow between the three species. Genomic landscapes of differentiation were correlated across pairwise comparisons, with relative divergence associated with local reductions in nucleotide diversity but mostly decoupled from absolute divergence. Peaks of differentiation were concentrated in putative pericentromeric regions, consistent with linked selection in shared low-recombination regions. In contrast, a large region of low relative differentiation on chromosome 2 showed elevated diversity across all species and overlapped a prominent cluster of chemosensory vomeronasal type-2 receptor (V2R) genes. Overall, our results suggest that shared demographic history and linked selection shape a common genomic landscape in *Hydrophis*, while chemosensory receptor gene evolution may have contributed to ecological diversification during the land-to-sea transition of these unique marine reptiles.

## Introduction

Increased genome sequencing of non-model, or emerging model, species has extended studies of adaptation and speciation across an array of systems, including flycatchers (Ellegren et al., 2012), swallows (Schield et al., 2025), cichlid fish (Kautt et al., 2020; Meier et al., 2017, 2018), *Heliconius* butterflies (Kronforst et al., 2013; Martin et al., 2013, 2016), poplar trees (Ma et al., 2018), and sunflowers (Andrew & Rieseberg, 2013; Renaut et al., 2013). These systems have provided valuable opportunities to observe lineages at the early stages of divergence, often at multiple points along the speciation continuum. In many such cases, genome-wide scans have identified local regions of pronounced genetic differentiation that stand out against a background of relatively low differentiation between diverging lineages. These so-called “islands of differentiation” can reflect various divergence processes depending on the selection regime, timing of gene flow, and underlying genomic architecture. Various models have been developed to identify and explain the formation of genomic islands, particularly using population genetic summary statistics of diversity and divergence.

Initial models invoked differential gene flow in elevating differentiation at loci involved in reproductive isolation (“speciation genes”) above a relatively homogeneous genetic background (Feder et al., 2012; Wu & Ting, 2004). Support for this is found if genomic islands show elevated differentiation for both relative (*F_ST_*) and absolute (*d_XY_*) measures of divergence, and low intraspecific diversity (*π*) (Han et al., 2017; Irwin et al., 2018). Alternative models explain heterogeneous patterns of differentiation without reference to reproductive isolation, suggesting that lineage sorting and selection can build high differentiation at some regions while low differentiation is maintained at other regions due to the sharing of ancestral polymorphisms (Cruickshank & Hahn, 2014; Noor & Bennett, 2009). Under this scenario, recent positive and/or background selection in allopatry are expected to generate high *F_ST_* and low *π,* but neutral *d_XY_*. Investigations of genomic islands characterised by high relative divergence and low intraspecific diversity have provided evidence of strong divergent positive selection in various taxa (Christmas et al., 2021; Liu et al., 2020; Schield et al., 2025). These studies have highlighted ecologically important phenotypes, such as mimicry wing patterns in *Heliconius* butterflies (Nadeau et al., 2012), as well as traits directly involved in reproductive isolation, including male plumage features in swallows (Schield et al., 2025).

Compared with positive and background selection, relatively few studies have highlighted the role of balancing selection in the formation of heterogeneous genomic landscapes. Evidence for balanced selection is found if genomic islands exhibit elevated *d_XY_*and high *π* but low *F_ST_*, reflecting the maintenance of advantageous polymorphisms among descendant lineages (Han et al., 2017; Irwin et al., 2018). This build-up and maintenance of adaptive genetic variation can result from several mechanisms, including heterozygote advantage, environmental and temporal heterogeneity in selection pressures, frequency-dependent selection, and antagonistic pleiotropy (where a genetic variant benefits one trait and is detrimental to another) (Hedrick, 2007). The importance of balancing selection is well established for immune genes, where maintaining multiple alleles can broaden defences against diverse pathogens (Croze et al., 2016). An analogous process has been proposed for chemosensory genes, such as olfactory receptors and their regulatory elements, the diversity of which underpins an ability to detect and discriminate odorants in various behaviours linked to survival and reproduction (Alonso et al., 2008; Liu et al., 2020). Chemosensory receptors, however, present an interpretive difficulty, because they are typically organised as tandem arrays and evolve under recurrent duplication, loss, and adaptive change (Nei & Rooney, 2005; Niimura & Nei, 2007). These complex dynamics of receptor gene family evolution can elevate local sequence heterogeneity and generate windowed summary statistics that resemble those expected under balancing selection. Hence, elevated diversity at chemosensory loci requires cautious interpretation.

All models of the formation of genomic islands emphasise the importance of genomic architecture and demography, particularly the timescale of divergence and history of gene flow. The effects of linked positive, balancing and background selection are promoted in regions with low recombination rates, especially those that also have high functional densities (i.e. harbour more targets for selection) (Seehausen et al., 2014; Wolf & Ellegren, 2017). This is seen particularly in systems that have retained stable patterns of recombination and gene density over successive speciation events; common genomic islands occur across multiple pairwise comparisons in these systems due to the long-term effects of linked selection in the same (low recombination) regions (Burri, 2017). This predicts that if a shared recombination landscape drives differentiation, the same genomic regions will be correlated across independent species pairs.

Here, we explore patterns of genomic differentiation and demographic history in an emerging system that is phylogenetically and ecologically distinct from previously studied taxa. The fully marine *Hydrophis* sea snakes share a common ancestor dated at only ∼1-8 million years old and exhibit the classic hallmarks of adaptive radiation (Kishida et al., 2025; Lee et al., 2016; Sanders, Lee, et al., 2013; Sanders, Rasmussen, et al., 2013). The ∼50 species in this clade show pronounced phenotypic diversity and occupation of novel morphospace (Sherratt et al., 2018, 2025), accompanied by high sympatric species richness across their Indo-Australian distribution (Voris & Voris, 1983). *Hydrophis* species have also undergone three-fold faster speciation compared to all other hydrophiine snakes, including the contemporaneous and geographically overlapping *Aipysurus-Emydocephalus* sea snakes (Lee et al., 2016; Sanders et al., 2010). Recently generated genomes show conserved chromosome-scale synteny, but widespread intra-chromosomal variations, and high repeat content in five *Hydrophis* species spanning the clade’s phylogeny (A. Li et al., 2021; Ludington et al., 2023).

The land-to-sea transition of these snakes was also accompanied by a major shift in chemosensory environment, from airborne to water-soluble cues. Squamates have few vomeronasal type-1 receptor genes but large vomeronasal type-2 receptor (V2R) repertoires, which detect dissolved chemical cues that are collected by tongue-flicking and delivered to the vomeronasal organ (Brykczynska et al., 2013; Schwenk, 1994). In snakes, V2R genes occur in long tandem arrays, with the largest reported on chromosome 2 (Hogan et al., 2021; A. Li et al., 2021). Chemosensory gene families are a plausible target of ecological diversification in *Hydrophis*, given that most species hunt concealed benthic prey and specialisation in prey type is a major axis of species differentiation in the clade (Kutsuma et al., 2018; Sherratt et al., 2018; Voris & Voris, 1983).

We re-sequenced single populations of three eco-morphologically distinct *Hydrophis* species that have overlapping distributions in northern Australia and Southeast Asia. *Hydrophis elegans* is a gracile, long bodied (up to 3m long) specialist on crevice and burrowing eels in depths up to 150m (C. Fry et al., 2001; Sherratt et al., 2018). *Hydrophis major* is of moderate length with a large head and strong jaws, and specialises on spiny toxic prey, especially plotosid catfish, in depths up to 50m (Shine et al., 2019). *Hydrophis curtus* is a short, thick-bodied generalist predator in shallow water (∼5-15m) and, unlike the other species, is predominantly day-active (Sherratt et al., 2018). The clear eco-morphological boundaries among these co-occurring species suggest near-complete or complete reproductive isolation, although recent phylogenomic analyses indicate a history of fluctuating population sizes, reticulate relationships, and extensive gene tree discordance, consistent with incomplete lineage sorting and possible interspecific gene flow during rapid speciation (Ludington et al., 2023; Ludington & Sanders, 2021).

Comparisons among these three species allow us to address three key questions. First, do the same genomic regions show elevated differentiation in every species pair, and do these regions share particular architectural features, as predicted under a shared recombination landscape? Second, do peaks of relative interspecific divergence (*F_ST_*) coincide with high absolute divergence (*d_XY_*), or simply reflect regions where diversity within species is reduced (*π*)? Finally, do chemosensory gene regions depart from genome-wide patterns in ways that are consistent with a role in ecological diversification?

## Methods

### Sample collection and sequencing

Samples from *Hydrophis major and H. elegans* species were collected and sampled during fieldwork in Western Australia, using procedures approved by The University of Adelaide’s Animal Ethics Committee (approval number S-2015-119 / 34903), under a fauna taking (scientific or other purposes) licence (regulation 25, number FO25000393) granted by the Department of Biodiversity, Conservation and Attractions of Western Australia. *H. curtus* were fished in the Gulf of Thailand and sourced from traders in wet markets in Cà Mau province in Vietnam. *Microcephalophis gracilis* was sourced by collaborators from commercial fisheries operating in coastal waters of the Emirate of Fujairah, United Arab Emirates. DNA was extracted from liver and tail tissues following the Gentra Puregene Tissue Kit protocol. DNA was sent to the South Australian Genomic Center (SAGC), where libraries were generated according to Illumina DNA Prep (M) guidelines (Part No. 10000000254 v10) and Illumina to MGI Library Conversion (MGIEasy Universal Library Conversion Kit, Part No. MGI000004155), and were sequenced on a MGI DNBSEQ-G400 (2×150bp paired-end). The *M. gracilis* sample was sequenced separately at the Ramaciotti Centre for Genomics. Libraries were prepped using Illumina DNA PCR-Free prep kit before being sequenced on an Illumina NovaSeq 6000 S4 flow cell (2×150bp paired-end). All sequence data were filtered for contaminants and low quality reads using *BBduk* v39.52 (*BBMap* – Bushnell B. – sourceforge.net/projects/bbmap), *Kraken2* v2.1.3 (Wood et al., 2019) and *Fastp* v0.23.2 (Chen, 2023). Sample accessions can be found in electronic supplementary table S1.

### Scaffolding *H. elegans* reference genome

To simplify downstream analyses, the *H. elegans* reference genome was scaffolded into pseudo-chromosomes using the genome-guided scaffolding module from *RagTag* v2.1.0 (Alonge et al., 2022). RagTag was run in *scaffold* mode (--remove-small), first aligning the *H. elegans* contigs to the *H. major* reference genome, orienting the contig sequences and finally filling gaps between adjacent contigs with *N*-sequences. This version of the *H. elegans* reference genome is used in subsequent analyses.

### Alignment and variant calling

#### Species matched reference

Whole genome resequencing datasets were aligned to their respective reference genomes (electronic supplementary table 1) for site frequency spectrum (SFS) and genome-wide within-species diversity analyses (see below). Data were aligned using *BWA2* (Vasimuddin et al., 2019), filtering for reads aligned in proper pairs with a minimum mapping quality of 20 using *samtools* v1.15.1 (--require-flags 3 –exclude-flags 4 –min-MQ 20) (Danecek et al., 2021). Duplicate alignments were marked by *sambamba* v0.8.2 (Tarasov et al., 2015) before depth statistics were calculated using *mosdepth* v0.3.3 (Pedersen & Quinlan, 2018). Joint genotyping was performed using *bcftools* v1.15.1 (Danecek et al., 2021), estimating genotype likelihoods with *mpileup* (-q 20-Q 20-a DP,AD) before calling variant and invariant sites using *call* (-m). Sites in close proximity to INDELs were removed in addition to the indels themselves (--SnpGap 10 --exclude-types indels), with variants then being normalised using *bcftools norm*. We then applied per-sample filters, setting variant genotypes to missing if read coverage dropped below 10x, or was greater than twice the sample’s average. Genotypes were also set to missing if allelic balance was < 0.2 or > 0.8. Variant sites were finally filtered on fraction missing (F_MISSING ≤ 0.2) within each species.

#### Common reference genome

Resequencing datasets were also mapped to the *Hydrophis major* reference genome, acting as the common coordinate system for all pairwise divergence analyses. The same alignment and variant calling pipelines from above were used, including the same variant filtering and missingness thresholds. Synteny analyses have indicated that the start of chromosome 2 is structurally complex (Ludington et al., 2023). To mitigate the impact of potential misassembly in the cross-species analyses, we masked variants that resided in problematic genomic loci. These regions were identified by mapping *H. major’s* PacBio HiFi reads back to the reference genome and flagging any region with haploid coverage as putative Z-chromosome misassembly.

### Demographic history

#### Estimation of the unfolded site frequency spectrum per-species

Unfolded site frequency spectra (SFS) were generated for each species from their reference-matched alignments using *ANGSD* v0.940 (Korneliussen et al., 2014). Ancestral reference genomes for each species were first created using *samtools consensus* (-a –min-MQ 20 –min-BQ 20 –show-ins no –show-del yes), substituting *M. gracilis* alleles into the reference sequences. Site allele frequency likelihood (SAF) files were then generated for each genome (-GL 1-doSaf 1-doCounts 1-minInd 5-setMinDepthInd 10-remove_bads 1-uniqueOnly 1-only_proper_pairs 1-minMapQ 30-minQ 30-sites species.sites), which were used as input to *realSFS* to create each species’ 1-dimension SFS (-bootstrap 50). Visualisation of SFS distributions, along with the calculation of expected allele frequencies under the neutral coalescent, were performed in R.

#### Effective population size inference

Effective population sizes through time were estimated for each *Hydrophis* species using the software *MSMC2* v2.1.1 (Schiffels & Wang, 2020) from their reference-matched alignments. Mappability files were created for each genome using *GenMap* v1.3.0 (Pockrandt et al., 2020) with a k-mer size of 49 and error allowance of 1 (-K 49-E 1). Variants were called per-sample/per-chromosome using *bcftools* mpileup v1.15.1 (-q 30-Q 30-C 50) and *call* (-c), and were converted into per-sample/per-chromosome VCF and mask files using *bamCaller.py*. The accessory script *generate_multihetsep.py* was then used to process these files, along with the genome mappability file, to create the final input files. *MSMC2* was run using the custom clock “1*4+1*3+25*2+1*4+1*6”, performing 100 EM-iterations (-i 100) with 20 bootstraps per sample. To scale the output, a mutation rate was estimated using the outgroup *M. gracilis* using a Jukes-Cantor corrected distance of *d =* 0.031, a split time of 12 million years (Lee et al., 2016), and a generation time of 5 years (Kishida et al., 2021), resulting in a per-site/per-generation mutation rate of μ = 6.45 x 10^-9^.

### Genomic divergence and introgression

#### Divergence statistics and correlation analyses

Relative (*F_ST_*) and absolute (*d_XY_*) measures of divergence were estimated between species pairs (*H. major - H. elegans, H. major - H. curtus* and *H. elegans* - *H. curtus)*, while nucleotide diversity (*π*) and Tajima’s *D* were estimated within each species. Fist, linkage decay plots were generated for each species using *popLDdecay* v3.42 (Zhang et al., 2019) to inform sliding-window size (electronic supplementary figure 1). Divergence statistics *F_ST_* and *d_XY_* were estimated for all pairwise species comparisons from the common-reference data in 100kbp non-overlapping windows using *pixy* v2.0.0.beta11 (*--stats dxy fst –window_size 100000*) (Korunes & Samuk, 2021). Genome-wide *F_ST_* was also calculated for each species comparison using *vcftools* v0.1.17 (Danecek et al., 2011). Within each species, nucleotide diversity (*π*) and *Tajima’s D* were estimated in 100kbp non-overlapping windows using *pixy* for both the reference-matched and common-reference datasets. The reference-matched data were used to obtain unbiased genome-wide estimates, while the common-reference results were used in genome-scans. Genomic windows with fewer than 10% of called sites were filtered out. Global estimates of *Tajima’s D* were also estimated from the SFS results using the R package *sfsr* (https://github.com/andrewparkermorgan/sfsr). *F_ST_* values were Z-transformed (*ZF_ST_*) to enable comparison of divergence landscapes between species pairs with different divergence times (Han et al., 2017). Windowed statistics were then correlated within each of the species-comparisons to explore their relationships, specifically: *F_ST_*-*d_XY_*, *π* - *F_ST_* and *π* - *d_XY_*. Additionally, windowed *F_ST_* and *d_XY_* measures were correlated between each of the pairwise species-comparisons to establish if shared or lineage-specific evolutionary processes were driving divergence across *Hydrophis*.

#### Genome-wide and sliding window introgression tests

Patterson’s *D* statistics (*ABBA-BABA* tests) were used to test for gene flow between the three *Hydrophis* species. *D* statistics use three ingroup populations and an outgroup with the relationship (((P1, P2), P3), O), testing for elevated shared variation between P2 and P3 compared to P1 and P3. The ASTRAL species tree reported in Ludington et al. (2023) was provided as the species topology to *Dsuite dtrios* v0.5r52 (Malinsky et al., 2021): *H. major* (P1), H. curtus (P2), H. elegans (P3) and *M. gracilis* (O). Introgression along the genome was also modelled using modified *ƒ*-statistics (*ƒ_d_*) in 100kb non-overlapping sliding windows via *ABBABABBAwindows.py* (https://github.com/simonhmartin/genomics_general; Martin et al., 2015). The same species tree was used, maintaining each window have at least 10% of sites called in addition to at least 200 SNPs. Estimates of *ƒ_d_* were correlated with windowed *d_XY_* estimates to explore the relationship between the signatures of divergence and introgression. As reduced *d_XY_* can artificially inflate estimates of *ƒ_d_* (Martin et al., 2015), we additionally calculated mean *d_XY_*between P2-P3 from *ƒ_d_* outlier windows (*ƒ_d_* ≥ 90^th^ percentile), expressing it as a percentage of mean *d_XY_*in the remaining background window. The expectation is that introgression should reduce absolute divergence, whereas ancestral structure can elevate *ƒ_d_*without a comparable reduction in *d_XY_*.

#### Detecting genomic outliers and assigning divergence profiles to genomic windows

Two separate approaches were used to examine genetic divergence between species pairs. First, high (*ZF_ST_*≥ 3) and low (Z*F_ST_* ≤-3) divergence windows were obtained, again only considering windows with genotype calls at 10% of sites. Permutation tests were implemented using regioneR (Gel et al., 2016) and used to test whether high-divergence windows had reduced absolute divergence (*d_XY_*) and nucleotide diversity (*π*; averaged between species) relative to windows in the genomic background. For each species comparison and summary statistic, the observed mean in high-divergence windows was compared to a null distribution generated via randomly resampling the same number of windows from all analysed windows (N = 10,000 permutations). Tests were one-sided (observed mean lower than the null) with significance assessed at α = 0.05.

As a second approach, divergence profiles were assigned to each 100kb window based on their *F_ST_* and *d_XY_* values. The four divergence profiles include allopatric speciation, balancing selection, recurrent selection and divergence-with-gene flow (Han et al., 2017; Irwin et al., 2018; Shang et al., 2023). Windows with *d_XY_* and *F_ST_* measures in the top one percent of values were classified as “divergence-with-gene-flow”. Windows with *F_ST_*values in the top one percent, but *d_XY_* values between the top and bottom one percent were classified as “allopatric selection”. Windows with *F_ST_* values in the top one percent but *d_XY_* values in the bottom one percent were classified as “recurrent selection”, an extension of “allopatric selection”. Finally, windows with *d_XY_* values in the top one percent and *F_ST_*in the bottom one percent were classified as “Balancing selection”. The location of centromeres was predicted using *Centrominer* v1.2.0 (Lin et al., 2023) to assist interpretation.

#### Annotation of vomeronasal type-2 receptor genes in *Hydrophis* snakes

Canonical vomeronasal type-2 (V2R) protein sequences were downloaded from Swiss-Prot, TrEMBL and NR protein databases (Sayers et al., 2022; The UniProt Consortium, 2025). To reduce redundancy, representative V2R sequences were obtained using *mmseqs2’s easy-cluster* workflow (Steinegger & Söding, 2017), using arguments –cov-mode 1 and –cluster-mode 2 to prevent short fragments from becoming representative proteins, whilst requiring that at least 90% of the target protein maps to the query (-c 0.9). Representative proteins were mapped to the *H. major* reference genome using *Miniprot* v0.13-r248 (H. Li, 2023), requiring at least 50% of the query protein align (--outc=0.5), resulting in candidate gene boundaries. V2R intron length distributions were generated from five snakes with RefSeq or GenBank gene annotations to inform alignment parameterisation (-G 45000) (electronic supplementary table 2). The resulting gene models were curated in IGV v2.17.4 (Thorvaldsdóttir et al., 2013), removing chimeric alignments that spanned multiple loci using *agat_sp_filter_feature_from_kill_list.pl* from AGAT v1.4.0 (Dainat, 2024). Protein alignment GFF3 files were then formatted using *agat_convert_sp_gxf2gxf.pl* (merge_loci = true). Protein sequences were extracted using *agat_sp_extract_sequences.pl*, and searched against the Pfam v37 database using *hmmscan* v3.4 (Eddy, 2011). Output files (*domtblout*) were filtered for 7-transmembrane (7tm) domains, requiring full alignment of the 7tm feature with c-evalues ≤ 1e^-5^. Sequences for 7tm domains were extracted using *bedtools getfasta* v2.31.1 (Quinlan & Hall, 2010), and clustered into representative sequence sets using CD-HIT v4.8.1 (-c 0.9) (W. Li & Godzik, 2006). Representative 7tm query sequences predicted from *H. major* were then searched against six sea snake genomes (electronic supplementary table 2) using *tBLASTn* v2.17.0 (Camacho et al., 2009), keeping only high-confidence complete alignments. Percentage identity distributions were generated from the alignments and used to inform filtering thresholds (*percentage identity* ≥ 60 used for all snakes), while alignments with overlapping coordinates were merged into single representative loci. Finally, candidate 7tm sequences from each snake were re-extracted and searched against Pfam to validate that the best hit for each sequence was in fact a 7tm domain, forming our final proxy V2R gene set.

## Results

### Sequence alignment and genotyping

Analyses in this study utilised two alignment datasets: species-matched and common-reference. Mapping rates after quality filtering ranged between 83 - 89% for species-matched datasets, dropping to 74.8 - 83% when mapping to the common *H. major* reference genome (excluding self-mapped *H. major* samples; electronic supplementary table S3). Average depth of coverage was similar between the species-matched and common reference genome datasets, differing on average by 2.1x depth-of-coverage (electronic supplementary table S3). The number of positions that could be genotyped was highest in the reference-matched datasets (electronic supplementary table S4-5), with fewer SNPs being called when samples were aligned to their respective genomes (electronic supplementary table S6). When calling variants against the common reference, joint-genotyping all samples resulted in the lowest number of variant and invariant sites being called, with the total number of genotyped sites increasing as the number of included samples decreased (electronic supplementary table S6). The *H. major* samples returned the lowest total number of genotyped sites, but had more variants than the other two species. Fewer positions being genotyped might be partially explained by the *H. major* samples having, on average, smaller sequencing libraries compared to the other samples (Supplementary figure 2).

### Effective population size through time

The demography of the three *Hydrophis* species was first investigated by comparing the unfolded site frequency spectrum (SFS) of each species to the derived allele frequencies expected under the standard neutral coalescent with constant population size (Fu, 1995). All three species of *Hydrophis* recorded elevated proportions of rare alleles (Figure 1A), which is consistent with population expansion, as the increase in effective population size (Ne) facilitates the mutation of new alleles (Watterson, 1975). The remaining derived allele count categories were typically close to, or slightly lower, than the expected proportions under the standard neutral model. Windowed and genome-wide estimates of Tajima’s *D*, which can also be used to detect departures from the expected SFS under neutral assumptions, also supported population expansion in all three species. Most genomic windows reported negative Tajima’s *D* values, while the genome-wide estimate from the SFS was negative in all three species, with negative values representing an excess of rare alleles in a population (Figure 1B; Table 1).

**Figure 1:**
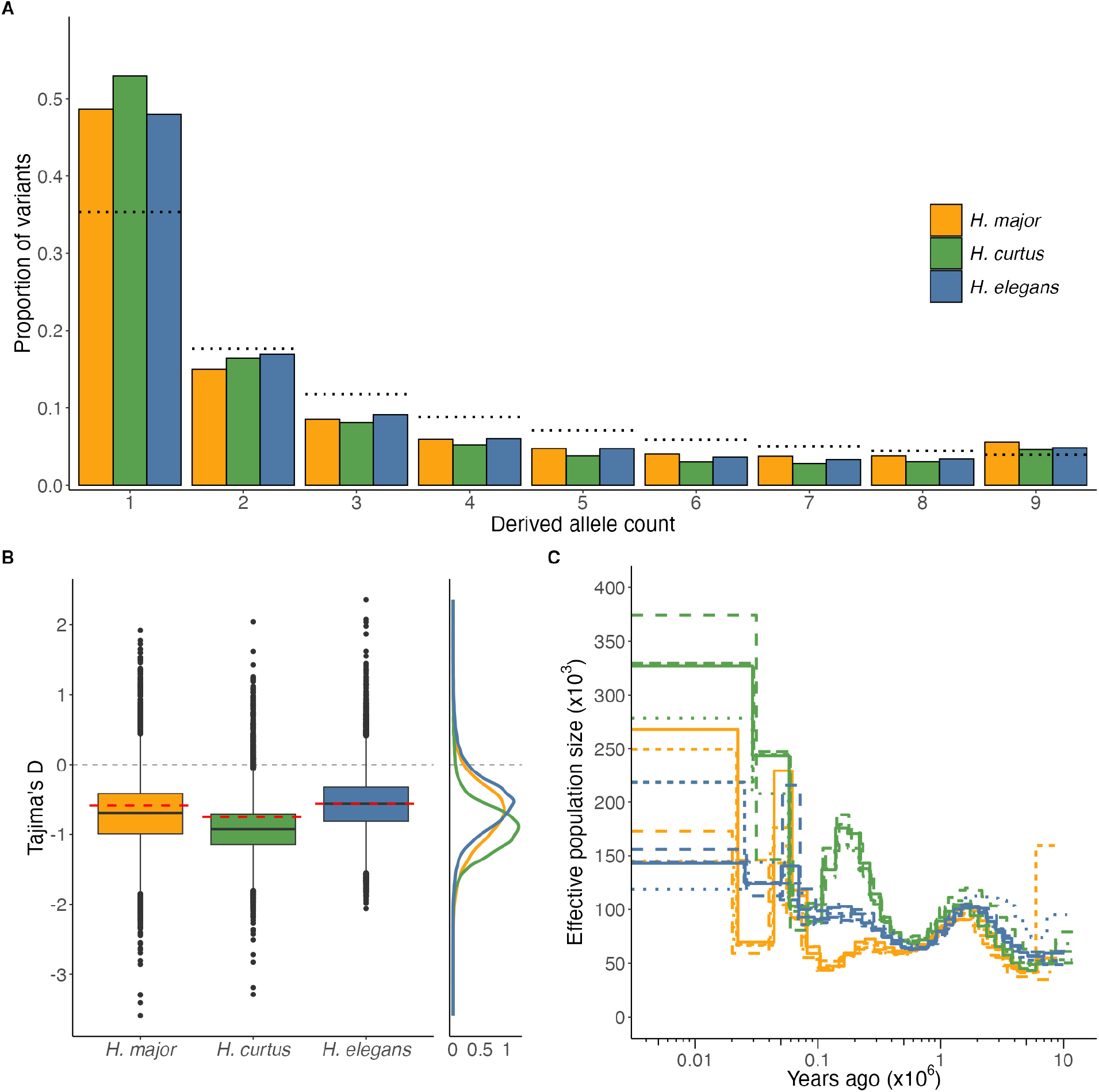
Demographic measures indicating recent population expansions in the three *Hydrophis* species. (A) Unfolded site frequency spectra for the three *Hydrophis* species. The dotted lines represent the expected variant proportions under the standard coalescent model with constant population size. (B) Boxplots of Tajima’s D calculated from 100-kb windows across autosomes, with the corresponding density plot of the windows shown to the right. Horizontal dashed lines in red represent genome-wide *Tajima’s D* values estimated from the SFS data. (C) MSMC2 plots of estimated effective population sizes (y-axis) through time (logarithmic x-axis), scaled using generation (5 years) and mutation rate (μ = 6.45 x 10^-9^ per-site/per-generation). Line-types represent different individuals within each species. A single *H. major* sample (KLS0631) had time indices 0-3 removed due to excessive Ne estimates (see electronic supplementary figure 3). Colours are consistent across each figure facet.

**Table 1:** Genome-wide average diversity/divergence statistics within and between species.

| Within species statistics |  |  | Between species statistics |  |  |  |
| --- | --- | --- | --- | --- | --- | --- |
| Species | Nucleotide diversity | Tajima's $D$ | Comparison | $d_{XY}$ | $F_{ST}$ (weighted) | $F_{ST}$ (unweighted) |
| *H. curtus* | 0.0024 | -0.75 | *H. curtus* - *H. elegans* | 0.0050 | 0.58 | 0.26 |
| *H. elegans* | 0.0020 | -0.56 | *H. curtus* - *H. major* | 0.0053 | 0.63 | 0.30 |
| *H. major* | 0.0016 | -0.59 | *H. elegans* - *H. major* | 0.0054 | 0.68 | 0.35 |

<sup>1</sup> Within species diversity statistics are calculated using reference-matched datasets.
<sup>2</sup> Between species diversity statistics are calculated from the masked common-reference datasets.

Fluctuations in effective population sizes (Ne) were also modelled using within-sample heterozygosity as an independent measure from the SFS (Martin et al., 2016). Historical estimates of each species’ Ne between 100mya and 5mya followed almost identical trajectories indicating a period of shared ancestry prior to a simultaneous divergence of the three lineages. Immediately following divergence, each lineage showed an upturn in Ne. Fluctuations in Ne were most pronounced in *H. curtus*, with the Southeast Asian lineage recording the highest Ne in the most recent time intervals. Of the two Australian lineages, *H. elegans* maintained the most stable population trajectory; this increased continually but was the lowest of the species in recent time intervals. *H. major* exhibited a shallow decline in Ne soon after its origin, followed by a steep rise and subsequent steep decline. In all species Ne was highest in the most recent time intervals (Figure 1C; electronic supplementary figure 3). These results support recent population expansions in the three *Hydrophis* species.

### Limited genomic evidence for historical introgression amid lineage discordance

Formal tests of gene flow between the three *Hydrophis* species returned little evidence of historical or ongoing introgression since speciation (figure 2A). Given the species tree from Ludington et al. (2023), genome-wide *D* failed to reach significance (*D* = 0.017, Z = 1.94, *P* = 0.052), reporting a low admixture proportion (*f_4_*-ratio = 0.86%). Although the majority site-pattern topology yielded a significant *D*-statistic (*D* = 0.058, Z = 10.5, *P* < 0.001), the corresponding *f_4_*-ratio remained low (2.3%), indicating a weak excess of allele sharing, consistent with the similar frequencies of the three site-patterns (30.4-35.4%). Notably, the concordant species topology (BBAA) was the least frequent pattern of the three (30.4%). Such similar frequencies indicate extensive incomplete lineage sorting, as expected for species that diverged near-simulataneously. Windowed estimates of introgression along the genome further supported the lack of allele sharing between the three species (figure 2B). Comparing estimates of *ƒ*_d_ and *d_XY_* in each window revealed a weak negative correlation between the two statistics (Spearman’s *ρ* =-0.37, *P* < 0.001), indicating measures of introgression (*ƒ*_d_) were typically elevated in genomic regions with lower absolute divergence. Whilst this pattern would be expected in the presence of gene flow, *ƒ*_d_ can also be inflated in windows of low *d_XY_* regardless of if there has been any historical or contemporary allele sharing (Martin et al., 2015). Introgression is expected to result in a relatively substantial reduction in absolute divergence in elevated *ƒ*_d_ regions. Mean P2-P3 *d_XY_* in high-*ƒ*_d_ windows as a percentage of the background mean was 78%. This modest reduction resembles the ancestral-structure and null simulations from Martin et al. (2015) more closely than their gene-flow simulations, though direct classification requires simulations parameterised for our study system.

**Figure 2:**
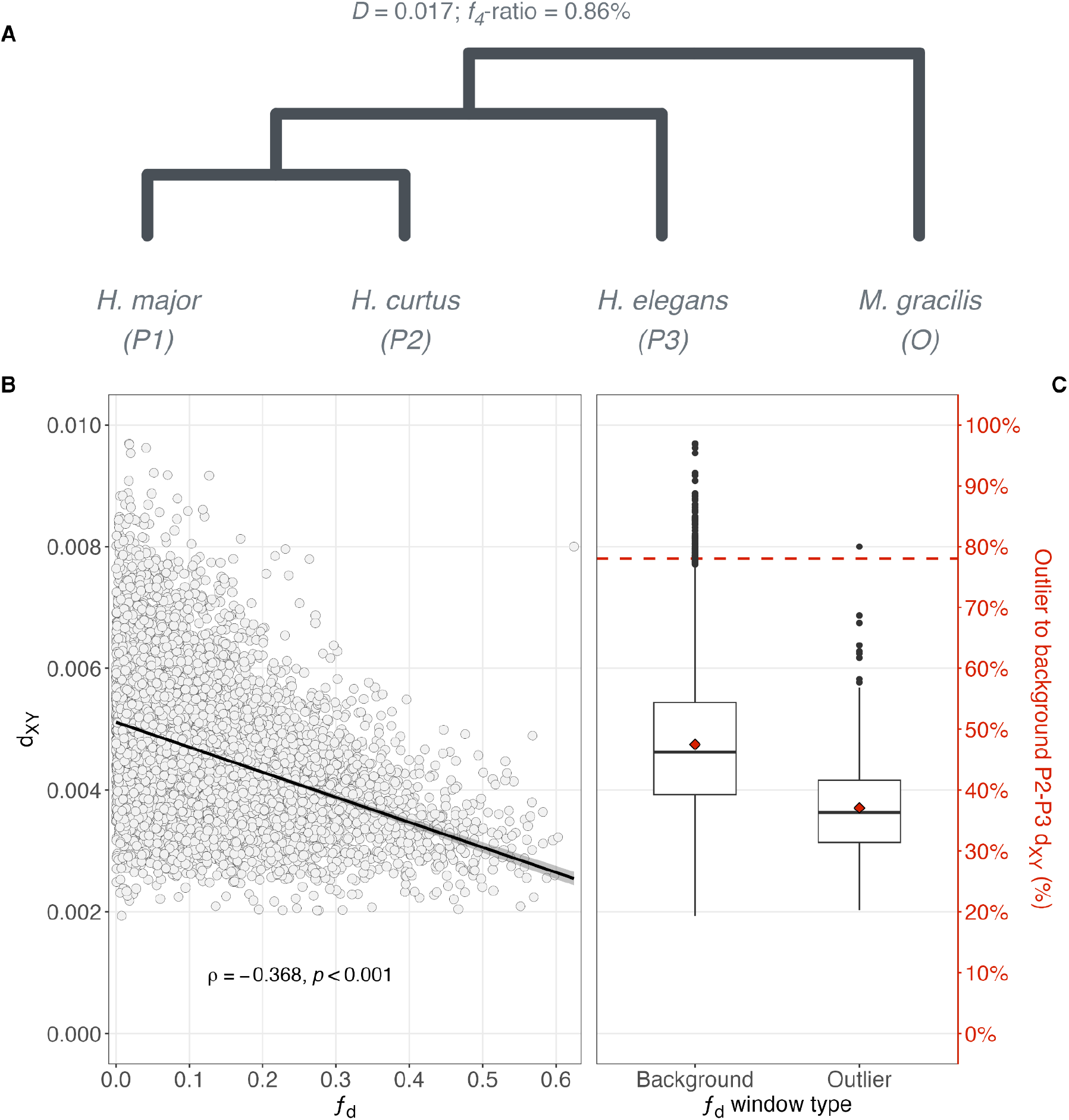
Limited support for introgression between the three tested species of *Hydrophis*. (A) Species tree from Ludington et al. (2023) used to polarise the genome-wide introgression test, with *H. major* as P1, *H. curtus* as P2, *H. elegans* as P3 and *M. gracilis* as the outgroup. The resulting *D*-statistic and *f_4_-*ratio estimates for this topology are shown above the tree. Branch lengths are not to scale. (B) Relationship between *ƒ_d_* and P2-P3 *d_XY_*across 100kb windows. Each point represents a 100kb window. The solid line shows the linear regression fit and the shaded band its 95% confidence interval. The correlation statistic is reported on the panel. (C) Distribution of P2-P3 *d_XY_* in high-*ƒ_d_* (outlier; ≥ 90^th^ percentile) and background 100kb windows. Red diamonds show the group means. The dashed red horizontal line shows the mean P2-P3 *d_XY_*of the outlier windows as a percentage of the background mean P2-P3 *d_XY_*. Facets B and C share the left y-axis. The red percentage line is interpreted against the red right y-axis.

### Genome-wide patterns of genetic diversity and divergence

Genome-wide diversity and divergence estimates are presented in table 1. Intraspecies diversity (*π*) ranged from 0.0016 to 0.0024, with absolute genetic divergence (*d_XY_*) ranging between 0.0050 to 0.0054, indicating the three focal species are genetically similar. In contrast, relative divergence (*F_ST_*) values were elevated between the three comparisons spanning 0.58 to 0.68, likely a product of the reduced nucleotide diversity. Average Tajima’s *D* was negative in all three species ranging from-0.75 to-0.56, signifying an excess of rare alleles in each species group. Correlations of windowed estimates of diversity and divergence statistics within each species revealed consistent patterns (figure 3). The distribution of windowed estimates of *π* were similar between species (figure 3A-C), and were negatively correlated with *F_ST_* with similar correlation coefficients across all comparisons (Spearman’s *ρ* =-0.637 to-0.697, *P* < 0.001) (figure 3D-F). Moderate significant positive correlations were observed between *π* and *d_XY_* in the three comparisons (figure 3G-I), again with similar correlation coefficients (Spearman’s *ρ* = 0.476 to 0.596, *P* < 0.001). Ostensibly zero correlation was observed between *F_ST_* and *d_XY_* (Spearman’s *ρ* = 0.0352 to 0.0643, *P* < 0.001), indicating that windows with elevated *F_ST_* are likely a product of low intraspecies diversity rather than elevated absolute divergence (figure 3J-L; electronic supplementary figure 4) (Holsinger & Weir, 2009). Pairwise correlations of windowed interspecies divergence statistics revealed that divergence profiles were largely consistent across the genome between each of the species comparisons, with both *F_ST_* (Spearman’s *ρ* = *0.672* to *0.693, P* < 0.001) and *d_XY_* (Spearman’s *ρ* = *0.713* to *0.788, P* < 0.001) showing significant moderate to strong positive correlations (electronic supplementary figure 5). While these pairwise comparisons are not evolutionarily independent, these results when taken together indicate that the genetic divergence between these three species has likely been shaped by shared evolutionary forces.

**Figure 3:**
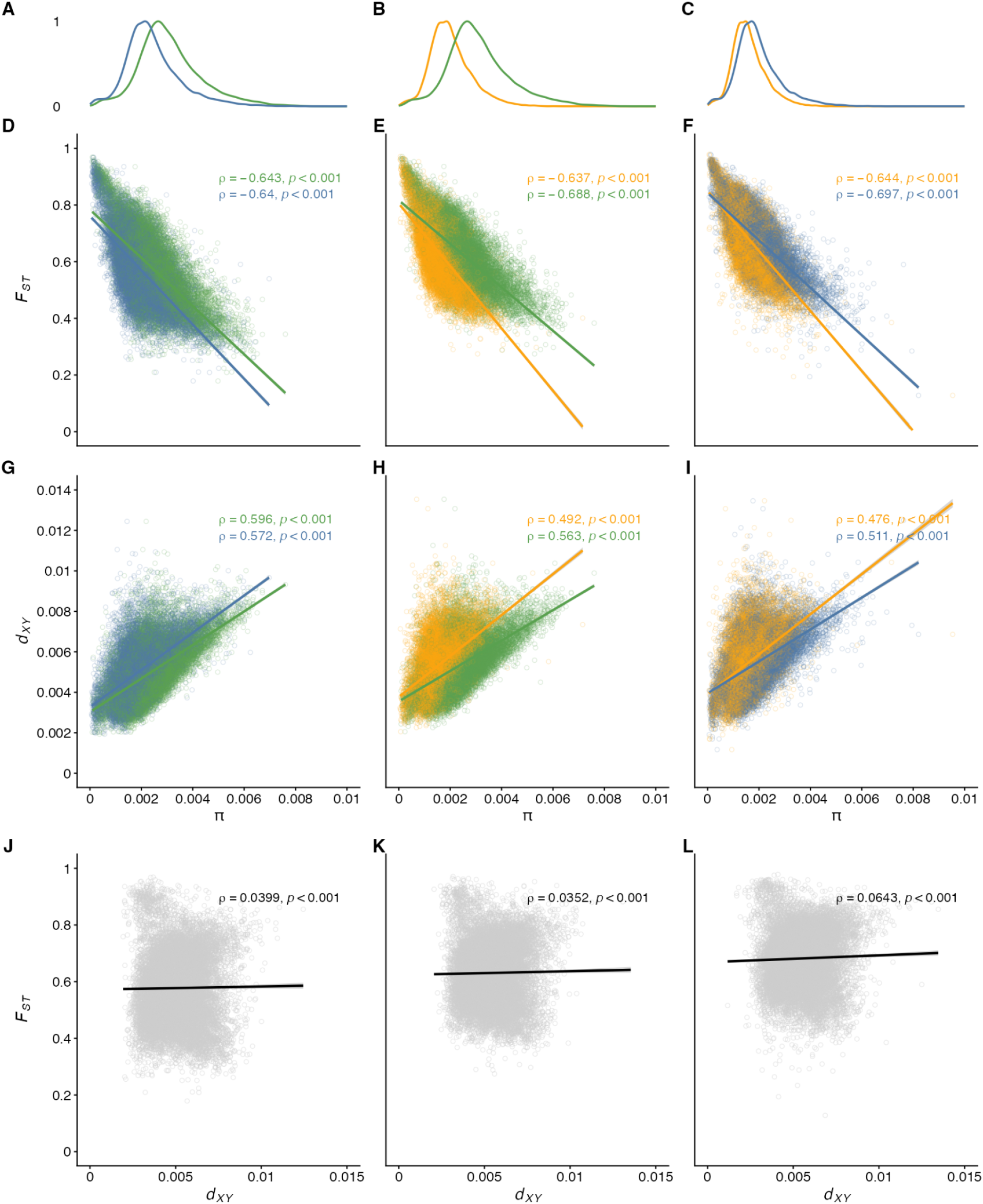
Relationships between within-species nucleotide diversity (*π*) and between-species relative and absolute divergence metrics (*F_ST_* and *d_XY_*) for all pairwise species comparisons. (A–C) Distributions of windowed *π* for species *H. elegans* (blue), *H. curtus* (green), and *H. major* (yellow). (D–F) Relationships between *π* and *F_ST_* for each pairwise species comparison. Within each panel, *π* from each species in the comparison is plotted against the same *F_ST_* values, with separate point clouds and regression lines. All comparisons show strong negative correlations, suggesting that genomic regions of reduced within-species diversity are driving the elevated relative divergence. (G–I) Relationships between *π* and *d_XY_* for the same species comparisons. Positive correlations in all species point to regions with higher nucleotide diversity also exhibiting higher absolute divergence, consistent with shared influences of mutation rate and/or recombination rate on both *π* and *d_XY_*. (J–L) Correlations between *d_XY_* and *F_ST_* for each comparison. Points represent the same genomic windows in each comparison. Very weak relationships (Spearman’s ρ < 0.07) indicate that variation in *F_ST_* is not driven by increases in *d_XY_*.

### Genome scans and divergence profiling

Genome scans and divergence profiling revealed similar patterns across all three species comparisons (figure 4; electronic supplementary figure 6). A total of 32, 34 and 2 windows were classified as high-divergence (*ZF_ST_* ≥ 3) in the *H. curtus - H. elegans*, *H. curtus - H. major* and *H. elegans* - *H. major* comparisons, resepectively (electronic supplementary table S7). In all comparisons, high-divergence windows clustered predominantly at putative centromeric loci (figure 4A-B), reporting significantly lower mean *d_XY_* and *average-π* in both the *H. curtus - H. elegans* and *H. curtus* - *H. major* comparisons relative to the genomic background (n = 32 and 34; all empirical *P* ≤ 0.001; N = 10,000 permutations) (electronic supplementary figures 7-8). Permutation tests were not carried out for the *H. elegans* - *H. major* comparison due to the limited number of high-divergence windows. In contrast, the *H. elegans* - *H. major* comparison recorded the most windows classified as low-divergence (*ZF_ST_* ≤-3; n = 49), with both the *H. curtus* - *H. elegans* (n = 6) and *H. curtus* - *H. major* (n = 9) comparisons reporting considerably fewer (electronic supplementary table S7). In the *H. elegans* - *H. major* comparison, low-divergence windows resided primarily at the start of chromosome 2 (figure 4B), while in the *H. curtus* - *H. elegans* and *H. curtus* - *H. major* comparisons, the majority of low-divergence windows were contained to a 9Mb region on chromosome 6 (chr6:58-67Mb).

**Figure 4:**
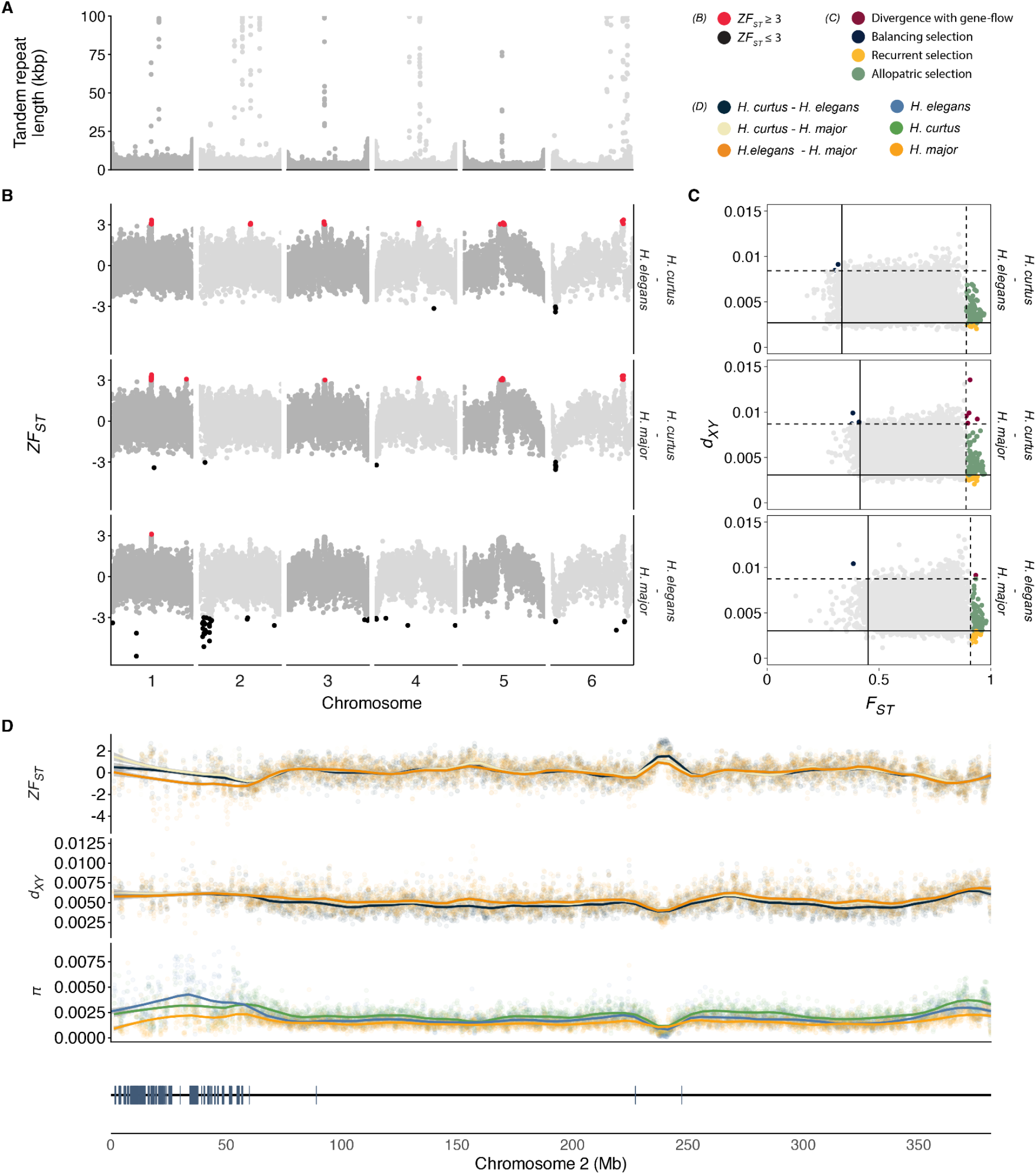
Genomic landscape of divergence in *Hydrophis* species. (A) Centromere predictions based on tandem-repeat (TR) content across the six macrochromosomes. Each point corresponds to a 100-kb window, with the y-axis indicating total TR length per-window. (B) Genome-wide scans of Z-transformed *F_ST_* (*ZF_ST_*) for each pairwise species comparison along the macrochromosomes in 100-kb windows. Windows with *ZF_ST_* ≥ 3 (elevated differentiation) and *ZF_ST_* ≤ –3 (reduced differentiation) are highlighted in red and black respectively, while grey points show the genome-wide distribution of all other windows. (C) Joint distribution of *F_ST_* and *d_XY_* across 100-kb windows for the same species comparisons. Dashed lines indicate the upper 1% thresholds for both metrics, and solid lines indicate the lower 1% thresholds. Points are colored by divergence-profile category: green (Allopatric: high *F_ST_*, intermediate *d_XY_*), yellow (Recurrent gene flow: high *F_ST_*, low *d_XY_*), dark blue (Balancing selection: low *F_ST_*, high *d_XY_*), and dark red (Divergence with gene flow: high *F_ST_*, high *d_XY_*). (D) Divergence statistics along chromosome two for all three species comparisons. The gene track represents the positions of putative V2R genes along the chromosome.

In addition to the genome scans, we extracted the top and bottom 1% of windows in each comparison and applied a divergence-profile framework (figure 4C). Across the three comparisons, between 169 to 171 genomic windows could be assigned to one of four main evolutionary scenarios (1.0% of all windows). Most windows were classed as “Allopatric selection” (elevated *F_ST_*, low *π* and average *d_XY_*), accounting for 143 to 153 (83.6% to 90.0%) of classified windows (electronic supplementary table S8). The next most common classification was “Recurrent selection” (elevated *F_ST_* and *π*, with reduced *d_XY_*), accounting for 15 to 22 (8.82% to 13.0%) of windows. Both allopatric and recurrent windows clustered primarily around centromeric loci (electronic supplementary figure 9). Windows classified as “Balancing selection” (elevated *π* and *d_XY_* and reduced *F_ST_*) and “Divergence with gene-flow” (elevated *F_ST_*and *d_XY_* and low *π*) were scarce, totalling between one to three windows (0.6% to 1.75%), and zero to five windows (up to 2.92%) respectively. Importantly, we treat discrete divergence profile labels as descriptive bins for windows in the extreme quantiles of *F_ST_* and *d_XY_* considered together, not as standalone evidence of particular selective mechanisms.

Focussing on the low-divergence signal at the start of chromosome 2 saw *ZF_ST_* between *H. elegans* - *H. major* trend lower than both species comparisons involving *H. curtus* (figure 4D). Little deviation in absolute divergence was observed across this region in the three comparisons compared to the genome-wide average, while *π* showed considerable variability. Consequently, the low *ZF_ST_* at the start of the chromosome appears to be a product of elevated genetic variation that is shared or is at similar frequencies in both the *H. elegans* and *H. major* species resulting in low relative divergence. Additionally, this region of chromosome 2 in sea snakes is dense with vomeronasal type-2 receptor (V2R) genes (A. Li et al., 2021). Using full-length 7-transmembrane (7tm) domains as proxies for V2R genes identified a total of 909 putative V2Rs in *H. major*, 430 of which were localised in this region of chromosome 2 (figure 4D; electronic supplementary tables 9-10). No windows classified as “balancing selection” along chromosome 2 in the *H. elegans - H. major* were also identified as low *ZF_ST_*. Chromosome 2 V2R totals in other snake genomes ranged between 48 to 524 genes, though no misassembly curation was performed (supplementary tables 9-10). Comparing V2R counts to the same chromosome scale terrestrial snakes from Ludington et al. (2023), *Hydrophis* snakes appear to have higher V2R counts than *Thamnophis elegans*, and similar numbers to *Crotalus adamanteus* (electronic supplementary table 9).

## Discussion

Population resequencing of three ecologically divergent *Hydrophis* species reveals a largely shared genome-wide signal of diversity and differentiation. Relative divergence is most strongly predicted by local reductions in nucleotide diversity, while absolute divergence shows little to no relationship to relative divergence, producing parallel divergence landscapes across all pairwise comparisons. Demographic inference from the site frequency spectrum and *MSMC2* supports recent population expansions in all three *Hydrophis*. Together with weak genome-wide signatures of non-treeness but localised evidence of allele sharing, these patterns are more easily reconciled with shared ancestral history and recurrent linked selection on a common recombination landscape than with widespread, species-specific barrier loci. High-*F_ST_* windows concentrate in pericentromeric regions, while a large low-*F_ST_* segment on chromosome 2 overlaps a dense array of chemosensory genes, highlighting genomic architecture and sensory multigene families as key factors shaping genomic differentiation in *Hydrophis* sea snakes.

### Historical demography and population size change

Demographic signals from the unfolded site frequency spectrum (SFS) and from *MSMC2* are broadly concordant in suggesting recent population expansions in *Hydrophis major*, *H. elegans* and *H. curtus*. All species converge on low effective population size at the deepest time intervals, consistent with shared ancestry before lineage-specific trajectories diverge (figure 1A). We interpret absolute timescales cautiously as they depend on mutation rate, generation time, and clock model, and because each analysis summarises population history from limited geographic sampling (single populations per species). Within those limits, *H. major* and *H. elegans* show similar recent Ne dynamics, compatible with overlapping northern Australian shelf distributions, whereas *H. curtus* attains the highest recent Ne estimates and the strongest temporal fluctuations, consistent with its broader Indo-West Pacific range and deeper phylogeographic structure relative to *H. major* and *H. elegans,* which are primarily endemic to the Australian region (Ukuwela et al., 2014).

Shallow marine habitats in this region have repeatedly gained and lost area through Plio-Pleistocene sea-level change (Cheng & Faidi, 2025; Voris, 2000), a setting commonly invoked to explain recent demographic expansions (and earlier bottlenecks) in coastal marine taxa (Ludt & Rocha, 2015). Such dynamics plausibly contribute to recent Ne increases in *Hydrophis*, although range shifts can also alter coalescent Ne estimates when formerly isolated demes reconnect through migration (Wang & Whitlock, 2003). The latter caveat may be especially relevant for *H. curtus*, for which fragmentation and reconnection of populations across the Indian and Pacific basins could interact with range fluctuations to shape Ne trajectories. Moving from qualitative agreement among SFS-based and pairwise-coalescent summaries to quantitative reconstruction of population dynamics and gene flow through time will require tests of region-specific demographic models and joint inference across multiple populations per species.

### Limited evidence for historical introgression

As *Hydrophis* is a rapid radiation, we formally investigated whether historical or contemporary gene flow has played a role in shaping the evolution of the three extant species. Genome-wide tests for introgression showed no clear signal of allele sharing given the published tree provided. Additionally, candidate regions along the genome with signals of past gene flow could not be distinguished from shared ancestral variation. We therefore find no clear evidence for substantial historical gene flow between these species. Nevertheless, the similar frequencies of the BBAA, ABBA and BABA site-patterns indicate weak support for any single branching order of the three lineages, with their genomes instead demonstrating extensive genealogical discordance. This ambiguity is perhaps not unexpected given the broader context of the system. Shallow divergence and rapid radiation, together with the demographic expansion inferred here, can all increase the retention of shared ancestral variation and generate patterns closely resembling incomplete lineage sorting (Pamilo & Nei, 1988). This compounds the difficulty of using summary introgression statistics to date or quantify admixture from single-population resequencing alone, and makes historical and contemporary gene flow inherently difficult to distinguish in these snakes.

### Genomic landscapes of differentiation

The three species comparisons in this study are not statistically independent, but the similarity in divergence profiles across the same genomic regions was nonetheless striking. Heterogeneity in relative and absolute divergence is largely shared across comparisons rather than specific to a single species pair. Correlations of diversity statistics within species comparisons illustrated patterns that conform to expectations under models in which elevated relative divergence reflects local reductions in diversity more than genome-wide increases in absolute divergence, resembling patterns summarised across systems using joint diversity frameworks (Han et al., 2017; Irwin et al., 2018; Shang et al., 2023).

The impact of reduced nucleotide diversity within each sea snake species was evident in the results from the genome scans, permutation tests and divergence profiling. In each species comparison, absolute divergence was lower in the high-divergence outlier windows compared to the genomic background, with outlier loci routinely overlapping stretches of the genome classified with “allopatric” or “recurrent” selection profiles. These signals are inconsistent with barrier loci expectations under sympatry, where both relative and absolute divergence are expected to be elevated (Cruickshank & Hahn, 2014; Ravinet et al., 2017). Instead, due to the proximity of the outlier regions to centromeres, the most parsimonious explanation for the observed peaks is likely background selection. Almost all high-divergence regions in this study were in or around centromeres, regions of the genome known for their low recombination rates (Noor & Bennett, 2009). Such regions typically report reduced nucleotide diversity (Nachman, 2002), with homogenisation of the underlying sequence occurring via background selection and/or genetic hitchhiking (Feder et al., 2012; Smith & Haigh, 1974). Comparable divergence patterns have been reported in other systems (Turner et al., 2005), with reanalysis of some published data similarly supporting the role of genomic architecture as the likely driver of the elevated relative divergence signal in centromeric regions (Cruickshank & Hahn, 2014). This highlights the importance of interpreting diversity and divergence summary statistics alongside functional genomic features when examining the roles of demographic history and genomic architecture in shaping patterns of differentiation and selection.

### Elevated diversity at chemosensory receptor genes on chromosome 2

The most conspicuous departure from the shared divergence landscape occurred at the start of chromosome 2, where a region of low relative differentiation coincided with elevated nucleotide diversity in all three comparisons. This region contains an array of 433 putative vomeronasal type-2 receptor (V2R) genes. Several observations suggest that this array is a functionally important feature of sea snake genomes. Large repertoires of vomeronasal type-2 receptor (V2R) genes have been reported in squamates (Brykczynska et al., 2013), with some of the most comprehensive gene arrays in snakes existing on chromosome 2. Snakes use tongue flicking to sample chemical cues from the environment and deliver them to the vomeronasal organ, where sensory neurons express vomeronasal receptors, primarily V2Rs (Schwenk, 1994; Silva & Antunes, 2017). In *Hydrophis*, where trophic specialisation is a major axis of ecological differentiation and many species forage on concealed benthic prey, chemically mediated prey detection and discrimination are expected to be central to fitness (Kutsuma et al., 2018; Sherratt et al., 2018; Voris & Voris, 1983). Chemosensory signaling is also relevant to reproduction, as sea snakes use close-range chemical cues during courtship and mate recognition (Shine, 2005).

This ecological context makes V2R loci plausible candidates of balancing selection; however, given that they are commonly organised as tandem arrays and evolve under birth-death dynamics (Nei & Rooney, 2005), excessive local diversity at these loci can arise via multiple avenues. Receptor-family expansion, turnover, and episodic adaptive change, particularly in extracellular ligand-interacting regions, can elevate sequence heterogeneity and create strong local effects on windowed population-genetic summaries (Hogan et al., 2021, 2025; Nei & Rooney, 2005; Niimura & Nei, 2007). We therefore avoid direct inference of balancing selection, taking a more conservative interpretation that elevated polymorphism reflects evolution in a dynamic chemosensory multigene region, potentially coupled to divergent ecological demands among lineages (Niimura & Nei, 2007). This interpretation is also consistent with expression-level complexity of vomeronasal systems, in which individual sensory neurons express one or a few receptor genes, and repertoire diversity is expected to shape the range and resolution of chemical cue discrimination (Malnic et al., 1999; Martini et al., 2001; Wynn et al., 2012).

A separate caveat concerns technical aspects of assembly quality. The number of V2R genes varied considerably between the examined *Hydrophis* snakes, and were comparable or slightly higher in number compared to the terrestrial snakes that were explored. Some of this variability is likely to be attributed to biological processes, but tandem arrays are notoriously difficult to assemble and the impact of assembly artifacts in these regions cannot be overlooked. Independent synteny analyses already illustrate structural complexity, or perhaps assembly issues, near the start of chromosome 2 in *Hydrophis* (Ludington et al., 2023), while broader evidence from reptiles supports repeat-mediated movement and amplification of vomeronasal receptor gene families (Zhu et al., 2026). Although candidate misassembled regions were masked before population-genomic analyses in this study, residual uncertainty remains. As such, the need for accurate, well curated reference genomes is imperative to resolve this complex gene family in *Hydrophis* snakes.

## Summary

Three co-occurring, ecologically distinct *Hydrophis* species that diverged within the last few million years share a very similar genomic landscape of divergence and diversity. Peaks of relative differentiation cluster near centromeres and coincide with low nucleotide diversity within species rather than high absolute divergence. This pattern is most consistent with linked selection acting on a shared recombination landscape. In contrast, an extended region of low relative differentiation on chromosome 2 shows elevated diversity across all species and overlaps a prominent cluster of chemosensory vomeronasal type-2 receptor genes. Since chemical cues are central to foraging and mating, chemosensory gene families are plausible targets of ecological selection during the land-to-sea transition of elapid snakes. Demographic analyses reveal recent population expansion in all three species, with little evidence of pervasive introgression. However, localised allele sharing is supported between the two sympatric Australian species, and may have contributed to the parallelism in their genomic landscapes.

## Author contributions

AJL and KLS conceived the research. AJL performed quantitative analyses. Both AJL and KLS interpreted results and drafted the manuscript.

## Acknowledgements

We thank Arne Rasmussen, Vinay Udyawer, Jenna Crowe-Riddell, James Nankivell, Isaac Rossetto and Nguyen Van Cao and Balazs Buzas for their help with sample collection. Data was processed on the Phoenix HpC at Adelaide University.

## Data availability

Raw sequence data are available through the NCBI SRA under bioprojects PRJNA1130540 and PRJNA984433. All code related to the processing and analysis of this data is available on GitHub: https://github.com/a-lud/hydrophis-popgen.

## Supplementary Figures

**Supplementary figure 1:**
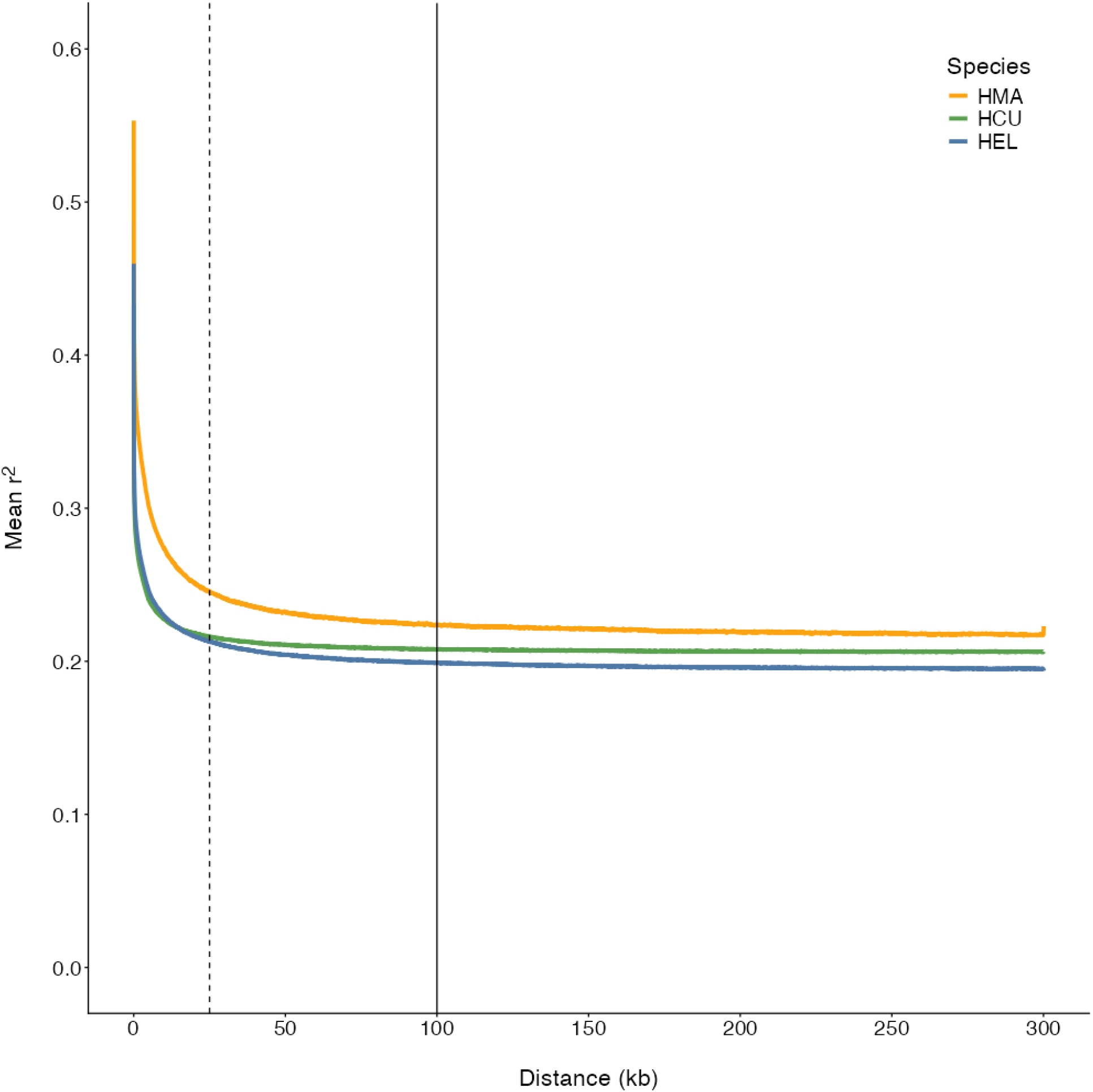
LD decay curves. Linkage decay curves estimated by *popLDdecay* from species-matched high-quality genetic variants. X-axis shows the genomic distance in kilobases, with the y-axis showing the mean *r^2^* value. Vertical dashed and solid lines show the 25kb and 100kb distances, respectively. Curves are predominantly flat by 100kb, but still show some decrease at 25kb.

**Supplementary figure 2:**
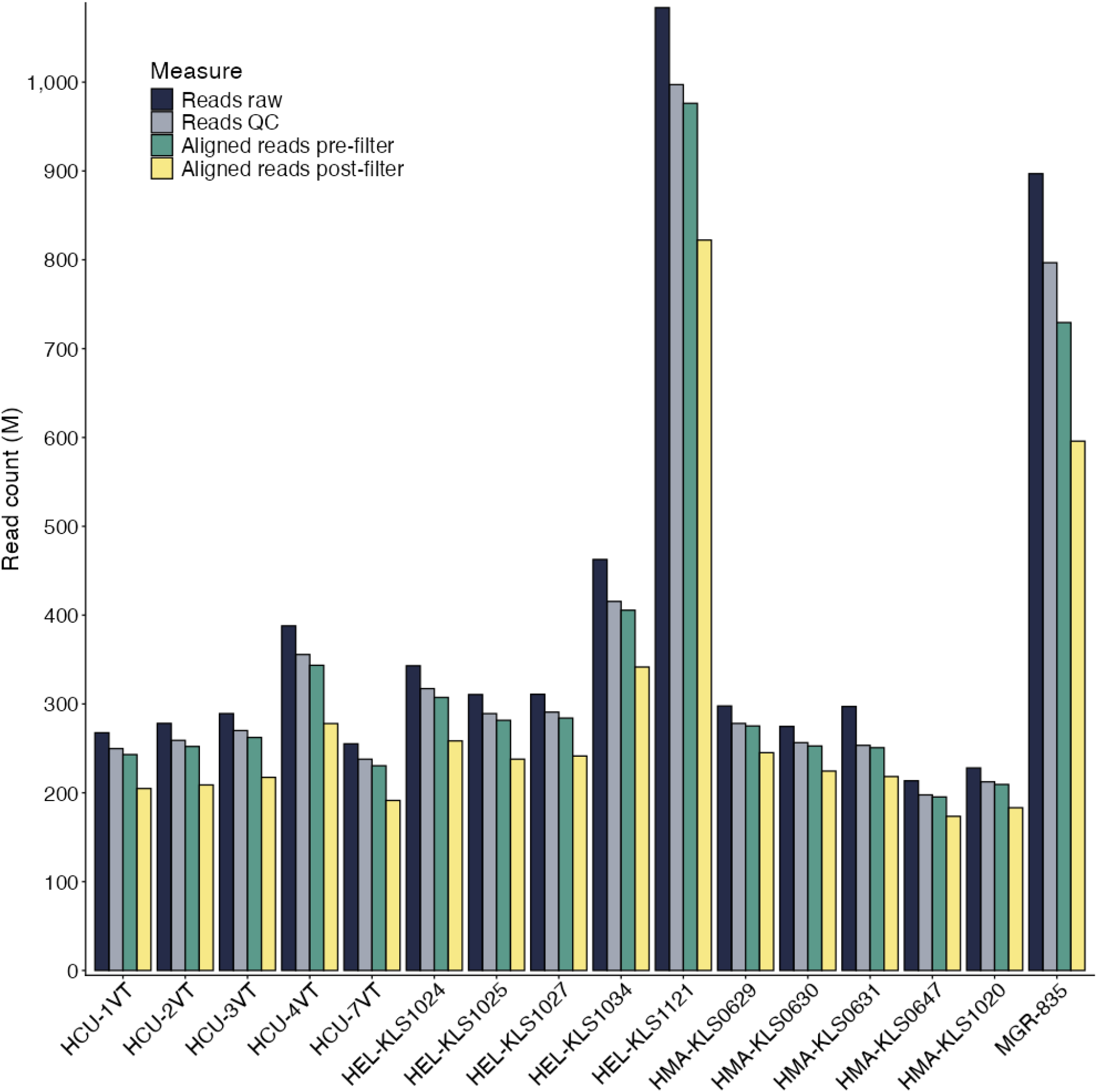
Alignment (common reference) Alignment statistics of all sixteen samples relative to the *H. major* reference genome. Columns are grouped by samples along the x-axis, with the number of reads (in millions) along the y-axis. Each coloured bar represents a different measure of the number of sequencing reads: black = raw total; grey = QC total; green = Aligned total; yellow = Aligned QC.

**Supplementary figure 3:**
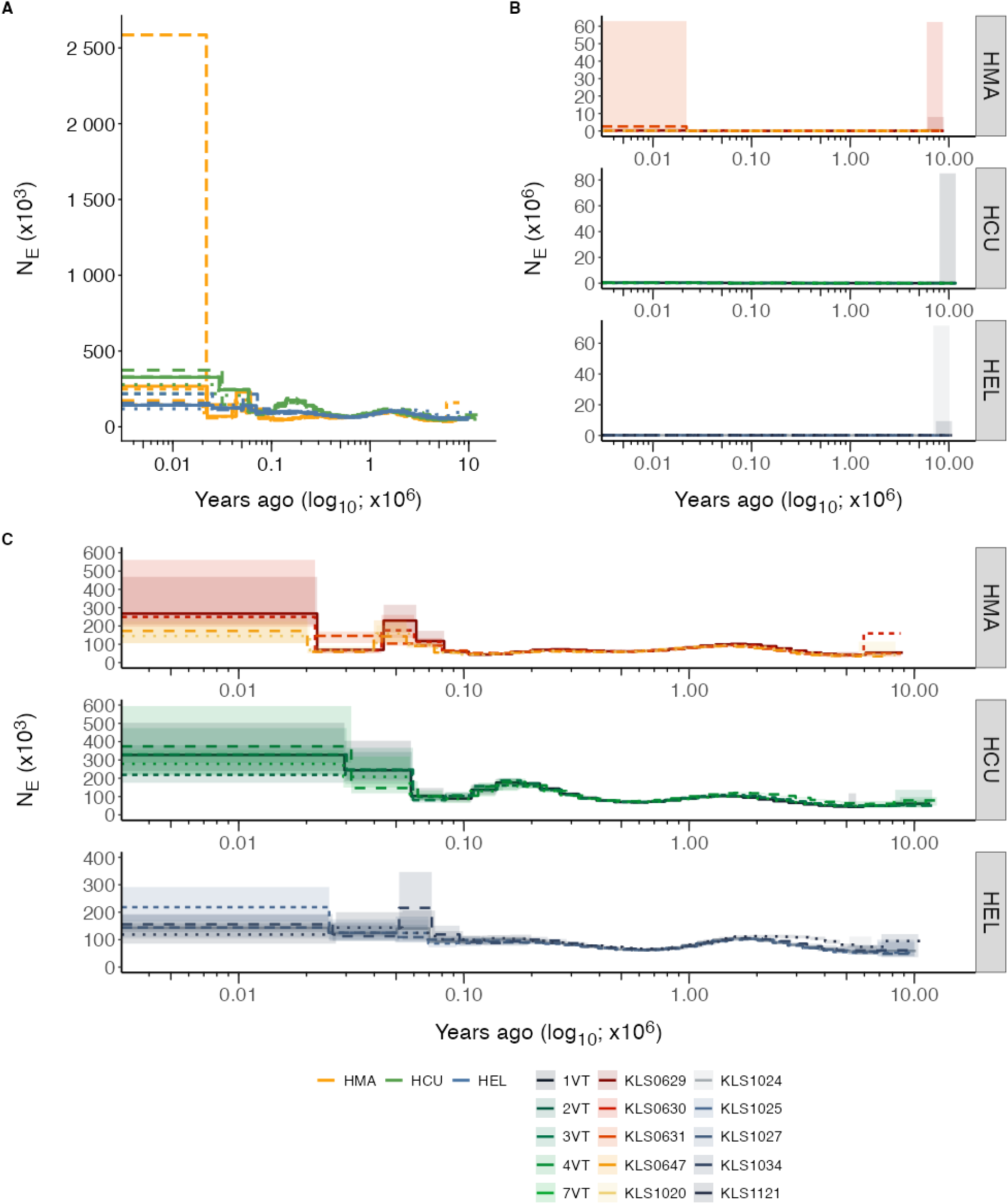
MSMC2 curves. MSMC2 was run independently on each biological sample, along with 20 independent bootstrap runs per-sample. A) Unfiltered *MSMC2* curves show a single *H. major* sample (KLS0631) with exceptionally large Ne estimates at the recent time intervals, but a curve matching the other *H. major* samples across all other time intervals. B) MSMC2 curves with bootstrap support shown as ribbons around the main MSMC2 curves from (A). Bootstrap estimates of Ne are exceptionally large in the distant past. C) Filtered MSMC2 curves with bootstrap support. Both primary MSMC2 curves and bootstrap estimates were filtered for maximum Ne ≤ 600,000, removing only spurious Ne estimates at the extreme time points, highlighting the tight bootstrap support at intermediate time-points. Different line-types are used to separate samples within species. Colours in facet A are independent of the shared colours in facets B and C. HMA = *H. major*, HCU = *H. curtus* and HEL = *H. elegans*.

**Supplementary figure 4:**
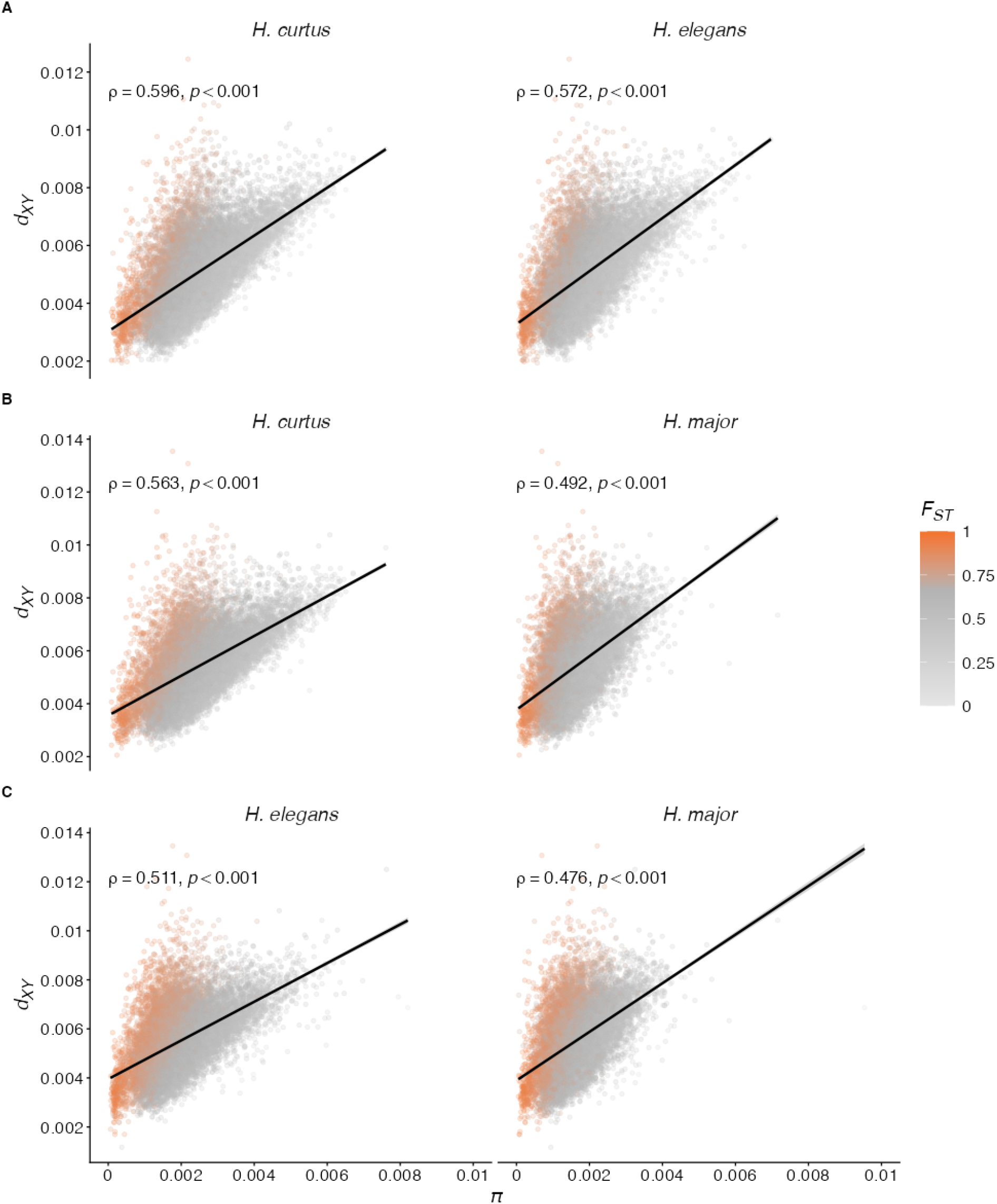
**Low-*π* drives elevated *F_ST_*** Windowed estimates of nucleotide diversity within species (*π*) correlated against absolute divergence (*d_XY_*) coloured by relative divergence (*F_ST_*). Colouring by *F_ST_*illustrates that instraspecific nucleotide diversity appears to be the driving force behind elevated relative divergence, not elevated absolute divergence. Columns in each facet represent the species for which the windowed *π* signal belongs, while each facet (row) represents the pair of species for which absolute divergence (*d_XY_*) was estimated. A) Comparison *H. curtus - H. elegans*. B) Comparison *H. curtus - H. major* and C) Comparison *H. elegans* - *H. major*.

**Supplementary figure 5:**
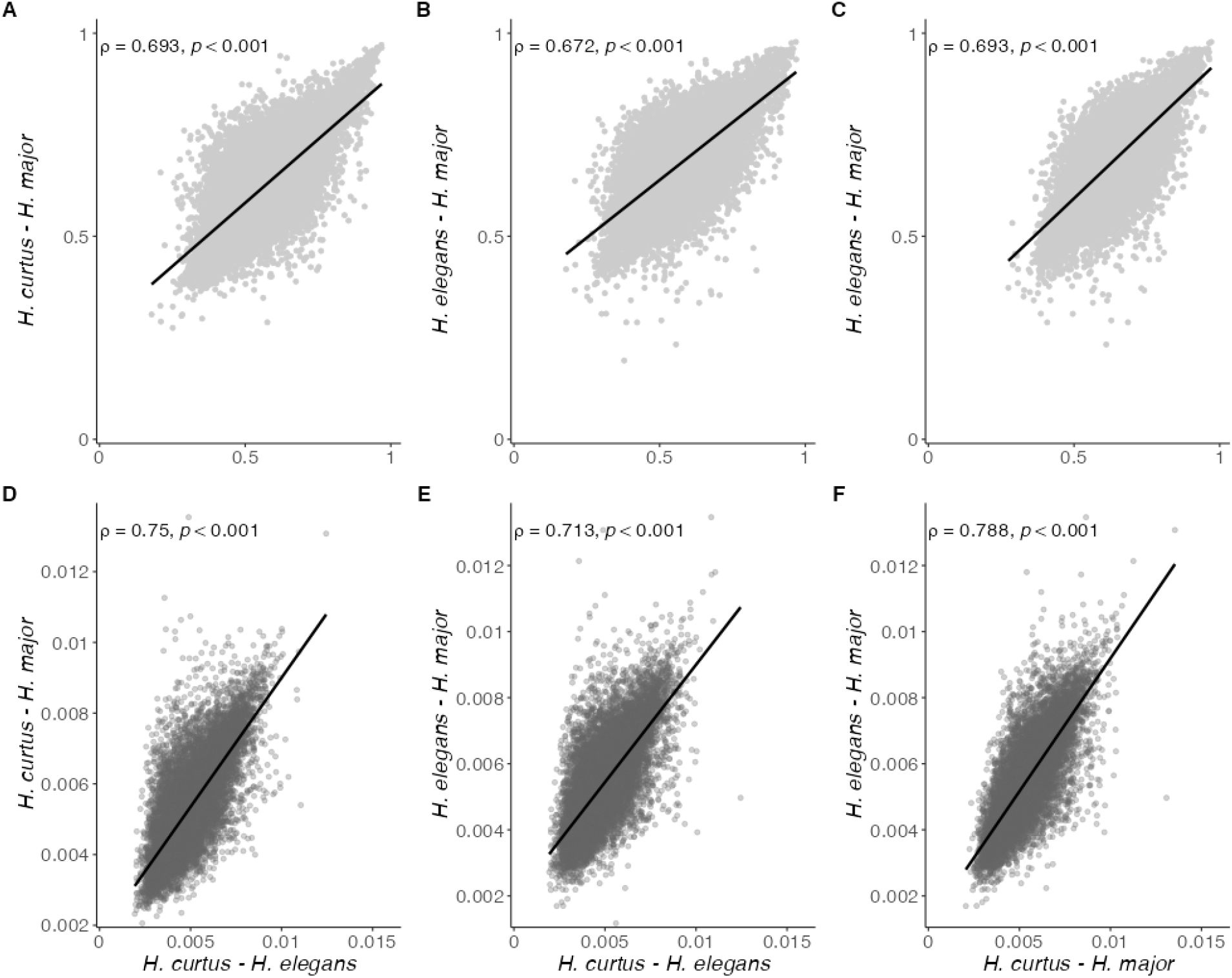
**Interspecies pairwise correlations of *F_ST_* and *d_XY_*** Spearman correlations of interspecies diversity statistics *F_ST_* (A-C) and *d_XY_* (D-F) between all three pairwise comparisons show moderate-to-high correlations. A-C) Correlation of the same genomic windows, and respective *F_ST_* estimates, between the three comparisons: *H. curtus - H. elegans* and *H. curtus* - *H. major* (A), *H. curtus - elegans* and *H. elegans - H. major* (B) and *H. curtus - H. major* and *H. elegans - H. major*. D-F) The same genomic windows and pairwise comparisons, but of *d_XY_*.

**Supplementary figure 6:**
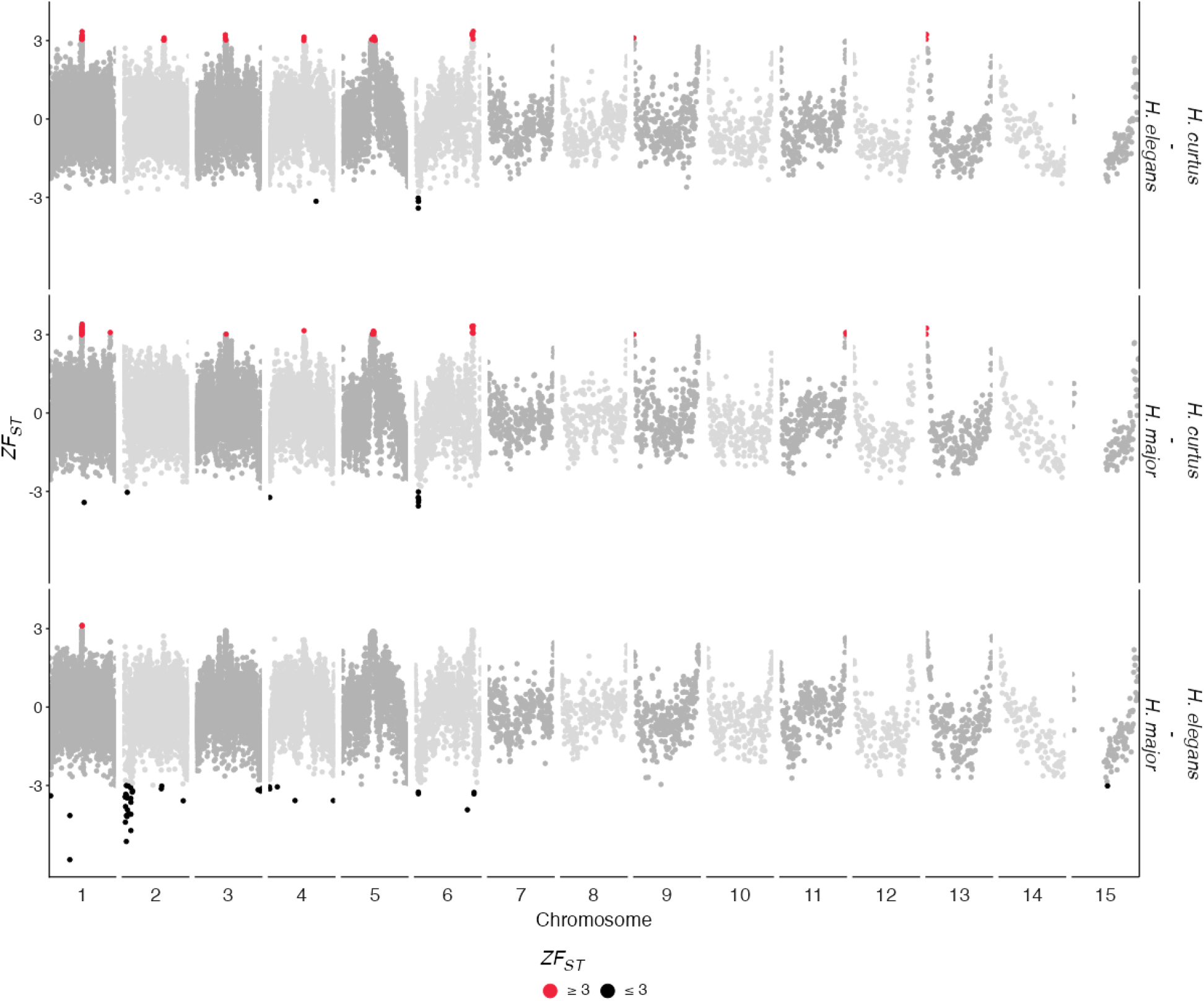
Genome-scan of divergence across the genome. Windows (100kb) of high-and low-divergence identified using *ZF_ST_* in each of the three species comparisons. Chromosomes are shown along the x-axis with *ZF_ST_* along the y-axis, with each of the three species comparisons represented by a facet. Windows coloured red represent high-divergence signals (*ZF_ST_* ≥ 3) while windows coloured black indicate genomic regions of low-diversity (*ZF_ST_* ≤-3). Diversity signals in the macrochromosomes are explored in the main-text figure 4. Diversity signals on the microchromosomes (chromosomes 7-15) are relatively sparse, with windows of high-divergence localised to chromosome starts, with only one low-divergence window reported on chromosome 15 in the *H. elegans* - *H. major* comparison.

**Supplementary figure 7:**
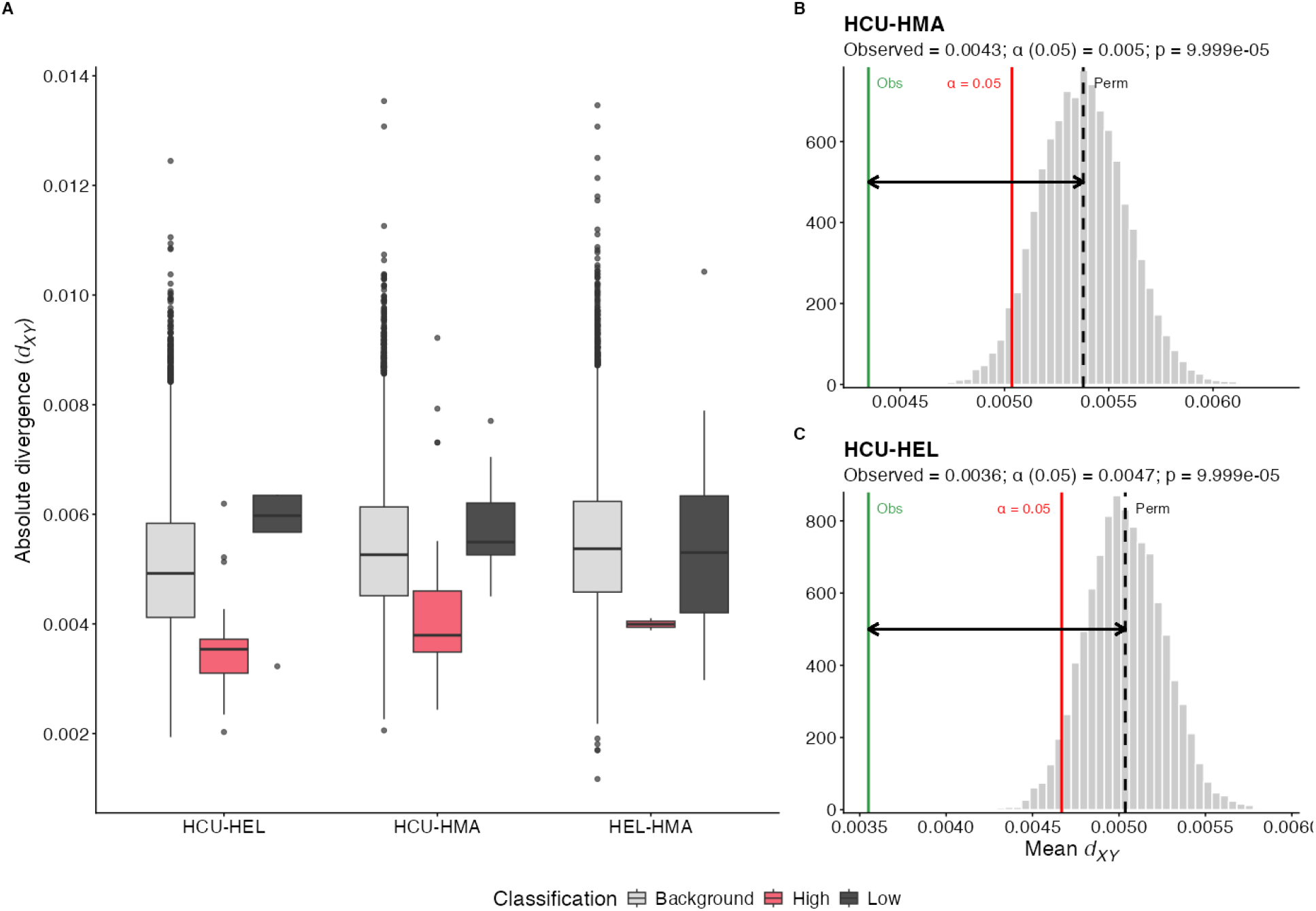
Variation in absolute divergence (*d_XY_*) and permutation results. Comparison of *d_XY_* between high-and low-divergence windows relative to the genomic background. A) Boxplots showing the distribution of *d_XY_* values in background (grey), high-divergence (red) and low-divergence (black) genomic windows. Each triplet of boxplots corresponds to a species comparison (x-axis). B-C) Permutation statistic results for *H. curtus - H. major* (B) and *H. curtus - H. elegans* (C). The black dashed line represents the null distribution (N = 10,000 permutations), the red line indicates the α = 0.05 threshold and the green line represents the observed mean.

**Supplementary figure 8:**
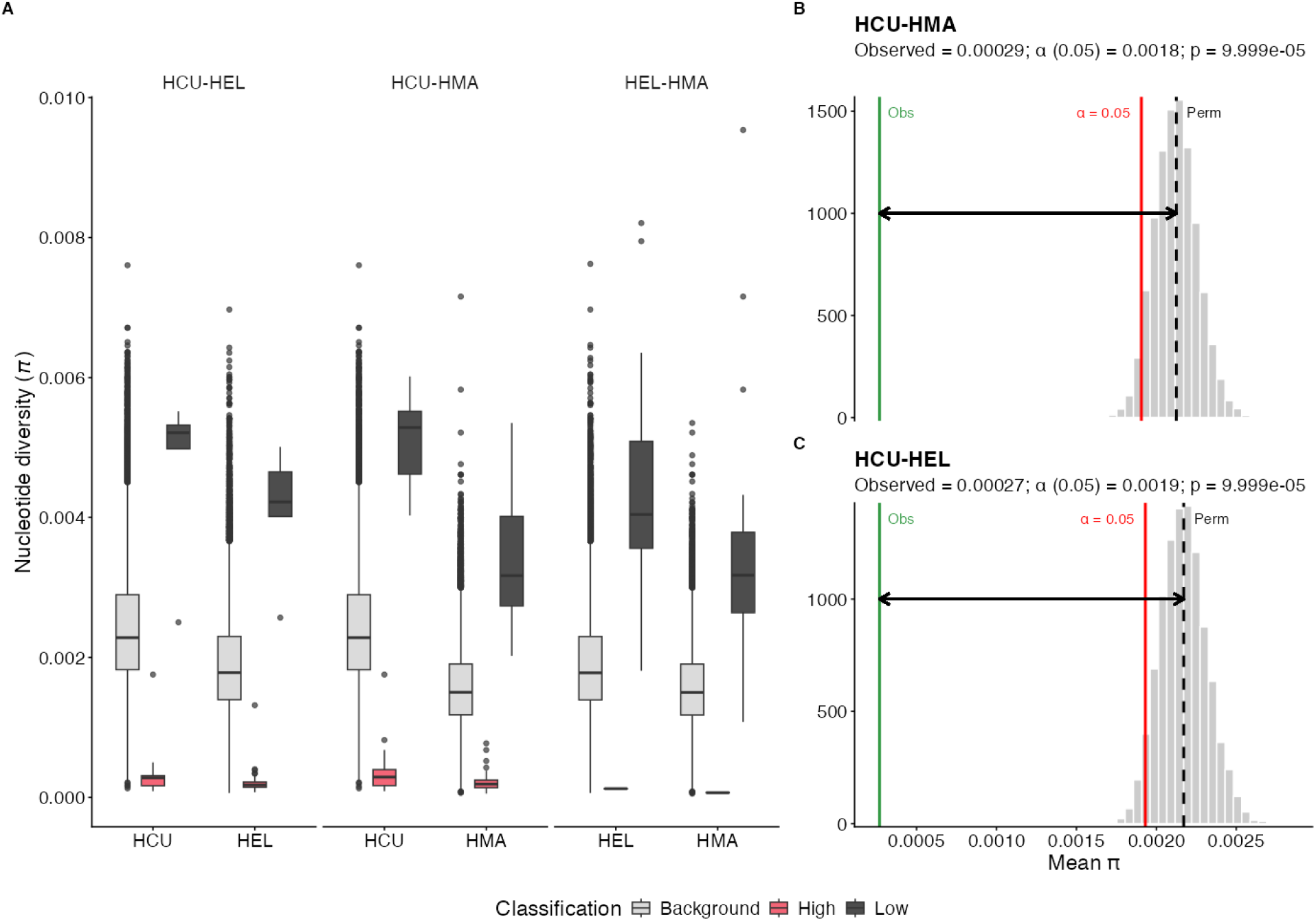
Variation in average nucleotide diversity (*average-π)* and permutation results. Comparison of *π* between high-and low-divergence windows relative to the genomic background. A) Boxplots showing the distribution of *π* values in background (grey), high-divergence (red) and low-divergence (black) genomic windows. Facets correspond to species comparisons, with each species forming a triplet of boxplots within the facet (x-axis). B-C) Permutation statistic results for *H. curtus - H. major* (B) and *H. curtus - H. elegans* (C). The black dashed line represents the null distribution (N = 10,000 permutations), the red line indicates the α = 0.05 threshold and the green line represents the observed mean.

**Supplementary figure 9:**
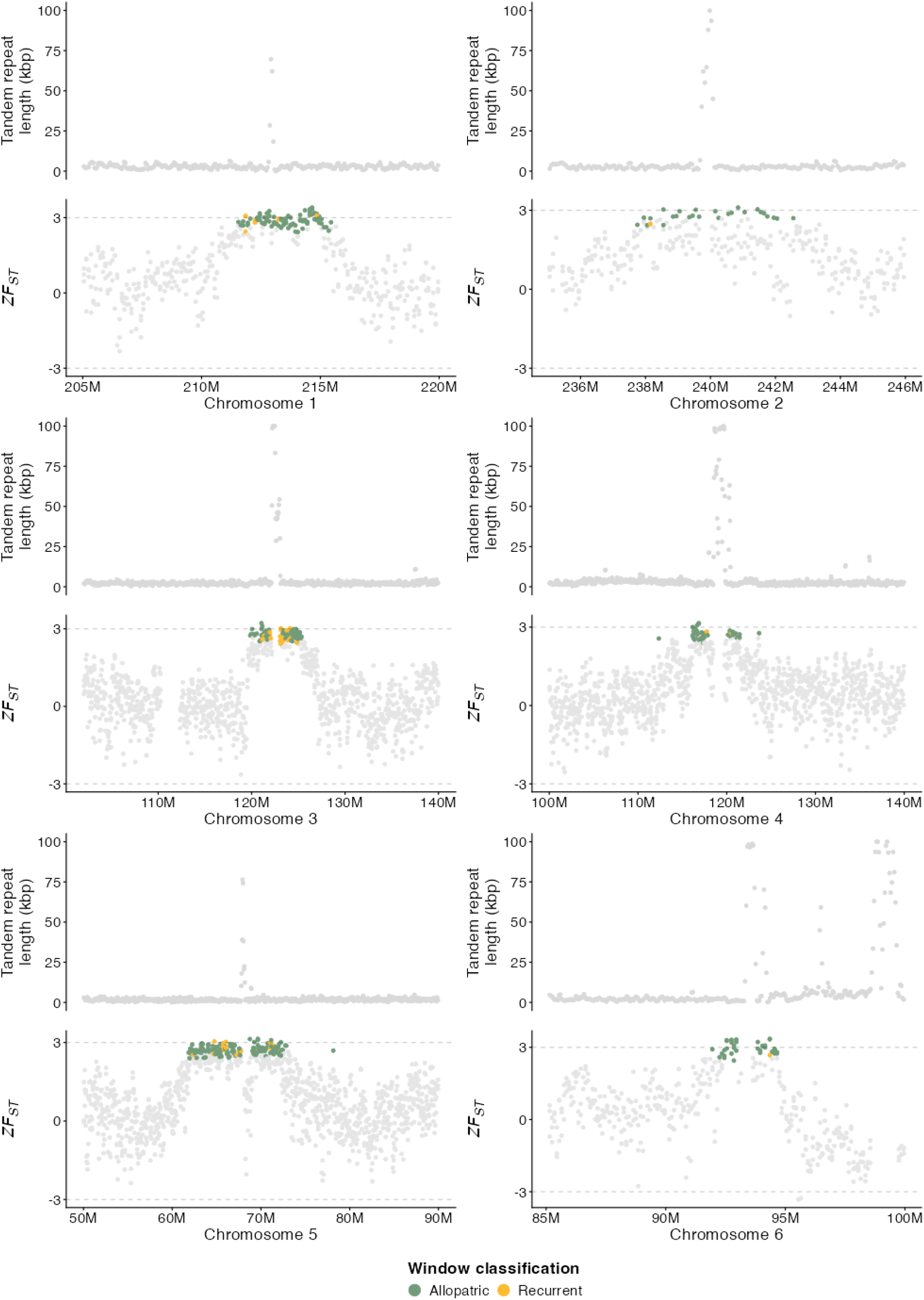
Putative centromeric regions enriched with windows profiled as “Allopatric” and “Recurrent”. Divenge profiling genome scan illustrates that many windows classified as “Allopatric” or “Recurrent” localise to genomic regions that are likely centromeric or pericentromeric. In each facet above, the top plot shows the tandem repeat content per-100kb window. The lower component shows *ZF_ST_* along the chromosome length, where each point is a 100kb window. Windows classified as “Allopatric” are in green, with “Recurrent” windows in yellow. Windows with no classification are in grey.

---

**Supplementary table 1:** Resequencing sample information.

| Supplementary table 1: Resequencing sample information |  |  |  |  |  |  | GenBank accession<br>(matched reference) |
| --- | --- | --- | --- | --- | --- | --- | --- |
| lab_id | species | locality | source | bioproject | biosample | run_accession |  |
| HCU-2VT | *Hydrophis curtus* | Gulf of Thailand | Purchased from Vietnam 2019 | PRJNA1130540 | SAMN42208568 | SRR29672487 | GCA_019473425.1 |
| HCU-3VT | *Hydrophis curtus* | Gulf of Thailand | Purchased from Vietnam 2019 | PRJNA1130540 | SAMN42208569 | SRR29672486 | GCA_019473425.1 |
| HCU-4VT | *Hydrophis curtus* | Gulf of Thailand | Purchased from Vietnam 2019 | PRJNA1130540 | SAMN42208570 | SRR29672485 | GCA_019473425.1 |
| HCU-7VT | *Hydrophis curtus* | Gulf of Thailand | Purchased from Vietnam 2019 | PRJNA1130540 | SAMN42208571 | SRR29672484 | GCA_019473425.1 |
| HCU-1VT | *Hydrophis curtus* | Gulf of Thailand | Purchased from Vietnam 2019 | PRJNA1130540 | SAMN42208567 | SRR29672488 | GCA_019473425.1 |
| HEL-KLS1024 | *Hydrophis elegans* | Broome, Australia | Field expedition 2018 | PRJNA1130540 | SAMN42208582 | SRR29672471 | GCA_033807725.1 |
| HEL-KLS1025 | *Hydrophis elegans* | Broome, Australia | Field expedition 2018 | PRJNA1130540 | SAMN42208583 | SRR29672470 | GCA_033807725.1 |
| HEL-KLS1027 | *Hydrophis elegans* | Broome, Australia | Field expedition 2018 | PRJNA1130540 | SAMN42208584 | SRR29672469 | GCA_033807725.1 |
| HEL-KLS1034 | *Hydrophis elegans* | Broome, Australia | Field expedition 2018 | PRJNA1130540 | SAMN42208585 | SRR29672468 | GCA_033807725.1 |
| HEL-KLS1121 | *Hydrophis elegans* | Broome, Australia | Field expedition 2021 | PRJNA984433 | SAMN35787919 | SRR24957355 | GCA_033807725.1 |
| HMA-KLS0629 | *Hydrophis major* | Broome, Australia | Field expedition 2015 | PRJNA1130540 | SAMN42208575 | SRR29672479 | GCA_033807585.1 |
| HMA-KLS0630 | *Hydrophis major* | Broome, Australia | Field expedition 2015 | PRJNA1130540 | SAMN42208576 | SRR29672478 | GCA_033807585.1 |
| HMA-KLS0631 | *Hydrophis major* | Broome, Australia | Field expedition 2015 | PRJNA1130540 | SAMN42208577 | SRR29672477 | GCA_033807585.1 |
| HMA-KLS0647 | *Hydrophis major* | Broome, Australia | Field expedition 2015 | PRJNA1130540 | SAMN42208578 | SRR29672476 | GCA_033807585.1 |
| HMA-KLS1020 | *Hydrophis major* | Broome, Australia | Field expedition 2018 | PRJNA1130540 | SAMN42208581 | SRR29672473 | GCA_033807585.1 |
| MGR-835 | *Microcephalophus gracilis* | Gulf of Oman | Field expedition 2021 | PRJNA1130540 | SAMN63240157 | SRR40747300 | N/A |

**Supplementary table 2:**
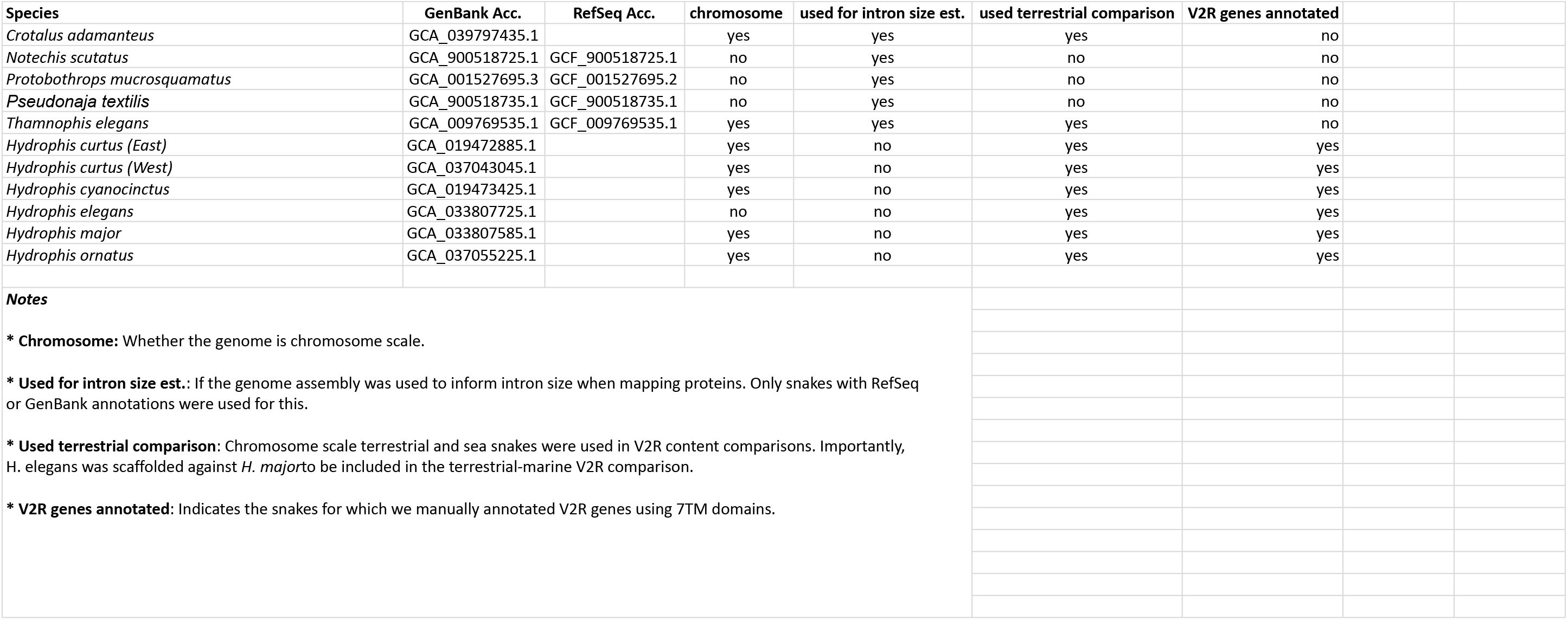
Snake genomes.

**Supplementary Table 3:** Sequence data and alignment summary statistics.

| Sample | Reads raw | Reads QC | Reads QC (%) | Reference matched |  |  |  |  | Common reference (*H. major*) |  |  |  |  |
| --- | --- | --- | --- | --- | --- | --- | --- | --- | --- | --- | --- | --- | --- |
|  |  |  |  | Aligned reads pre-filter | Aligned reads pre-filter (%) | Aligned reads post-filter | Aligned reads post-filter (%) | Mean coverage | Aligned reads pre-filter | Aligned reads pre-filter (%) | Aligned reads post-filter | Aligned reads post-filter (%) | Mean coverage |
| HCU-1VT | 267,586,314 | 249,808,162 | 93.36 | 245,724,658 | 91.83 | 216,617,855 | 80.95 | 16.5 | 243,036,206 | 90.83 | 204,861,542 | 76.56 | 15.19 |
| HCU-2VT | 278,164,778 | 259,019,694 | 93.12 | 254,348,152 | 91.44 | 220,737,699 | 79.36 | 16.73 | 252,184,260 | 90.66 | 208,856,404 | 75.08 | 15.42 |
| HCU-3VT | 289,188,448 | 270,066,406 | 93.39 | 264,789,746 | 91.56 | 229,733,049 | 79.44 | 17.42 | 262,324,182 | 90.71 | 217,401,099 | 75.18 | 16.04 |
| HCU-4VT | 387,932,328 | 355,716,264 | 91.70 | 346,730,436 | 89.38 | 295,071,532 | 76.06 | 21.92 | 343,548,386 | 88.56 | 277,923,735 | 71.64 | 20.09 |
| HCU-7VT | 255,169,496 | 237,797,988 | 93.19 | 232,877,686 | 91.26 | 202,636,596 | 79.41 | 15.4 | 230,433,330 | 90.31 | 191,401,176 | 75.01 | 14.15 |
| HEL-KLS1024 | 343,009,608 | 317,305,754 | 92.51 | 310,799,158 | 90.61 | 276,194,777 | 80.52 | 20.84 | 307,360,310 | 89.61 | 258,469,765 | 75.35 | 19.03 |
| HEL-KLS1025 | 310,601,808 | 289,153,862 | 93.09 | 285,036,352 | 91.77 | 254,553,438 | 81.95 | 19.26 | 281,608,700 | 90.67 | 237,847,020 | 76.58 | 17.57 |
| HEL-KLS1027 | 310,971,320 | 290,825,834 | 93.52 | 287,746,526 | 92.53 | 257,840,045 | 82.91 | 19.54 | 284,105,446 | 91.36 | 241,426,349 | 77.64 | 17.86 |
| HEL-KLS1034 | 462,669,448 | 415,463,134 | 89.80 | 410,638,262 | 88.75 | 364,921,124 | 78.87 | 27.45 | 405,580,174 | 87.66 | 341,604,172 | 73.83 | 25.09 |
| HEL-KLS1121 | 1,083,815,782 | 997,195,570 | 92.01 | 988,758,952 | 91.23 | 887,314,711 | 81.87 | 65.78 | 976,132,088 | 90.06 | 822,225,177 | 75.86 | 59.48 |
| HMA-KLS0629 | 297,733,702 | 278,099,610 | 93.41 | 275,213,204 | 92.44 | 245,153,649 | 82.34 | 18.25 | 275,213,204 | 92.44 | 245,153,649 | 82.34 | 18.25 |
| HMA-KLS0630 | 274,748,352 | 256,435,026 | 93.33 | 252,676,796 | 91.97 | 224,462,851 | 81.70 | 16.71 | 252,676,796 | 91.97 | 224,462,851 | 81.70 | 16.71 |
| HMA-KLS0631 | 297,168,064 | 253,449,224 | 85.29 | 250,803,256 | 84.40 | 218,289,061 | 73.46 | 16 | 250,803,256 | 84.40 | 218,289,061 | 73.46 | 16 |
| HMA-KLS0647 | 213,580,724 | 197,635,582 | 92.53 | 195,328,190 | 91.45 | 173,540,058 | 81.25 | 12.92 | 195,328,190 | 91.45 | 173,540,058 | 81.25 | 12.92 |
| HMA-KLS1020 | 227,979,122 | 212,416,560 | 93.17 | 209,411,136 | 91.86 | 183,187,176 | 80.35 | 13.63 | 209,411,136 | 91.86 | 183,187,176 | 80.35 | 13.63 |
| MGR-835 | 896,979,132 | 796,714,316 | 88.82 |  |  |  |  |  | 729,373,332 | 81.31 | 595,842,321 | 66.43 | 40.74 |
| **Notes** |  |  |  |  |  |  |  |  |  |  |  |  |  |
| * All percentages are calculated relative to the "raw" totals to represent total data loss relative to the starting amounts. |  |  |  |  |  |  |  |  |  |  |  |  |  |
| * Matched data: Samples had their data mapped to their respective reference genomes |  |  |  |  |  |  |  |  |  |  |  |  |  |
| - HCU = *H. curtus* reference genome |  |  |  |  |  |  |  |  |  |  |  |  |  |
| - HEL = *H. elegans* reference genome |  |  |  |  |  |  |  |  |  |  |  |  |  |
| - HMA = *H. major* reference genome |  |  |  |  |  |  |  |  |  |  |  |  |  |
| * Common reference: Samples had their sequence data aligned to the *H. major* reference genome. |  |  |  |  |  |  |  |  |  |  |  |  |  |

**Supplementary table 4:** Callable sites (matched reference genome)

| Supplementary table 4: Callable sites (matched reference genome) |  |  |  |  |  |  |  |  |  |  |  |
| --- | --- | --- | --- | --- | --- | --- | --- | --- | --- | --- | --- |
| *H. curtus* |  |  |  | *H. elegans* |  |  |  | *H. major* |  |  |  |
| sample | chr | length | called | sample | chr | length | called | sample | chr | length | called |
| *H. curtus* | chr1 | 438,509,360 | 335,452,060 | *H. elegans* | chr1 | 439,018,779 | 373,699,305 | *H. major* | chr1 | 434,526,931 | 311,599,672 |
| *H. curtus* | chr2 | 364,343,143 | 264,524,076 | *H. elegans* | chr2 | 375,143,852 | 301,128,589 | *H. major* | chr2 | 381,225,800 | 246,793,452 |
| *H. curtus* | chr3 | 266,229,956 | 202,566,842 | *H. elegans* | chr3 | 267,871,518 | 225,772,977 | *H. major* | chr3 | 268,276,564 | 184,940,572 |
| *H. curtus* | chr4 | 220,577,908 | 164,724,490 | *H. elegans* | chr4 | 220,543,031 | 185,027,479 | *H. major* | chr4 | 218,045,602 | 152,746,588 |
| *H. curtus* | chr5 | 145,831,796 | 106,270,208 | *H. elegans* | chr5 | 146,002,398 | 121,197,545 | *H. major* | chr5 | 142,905,876 | 97,378,071 |
| *H. curtus* | chr6 | 103,347,851 | 73,655,198 | *H. elegans* | chr6 | 106,234,579 | 85,269,245 | *H. major* | chr6 | 106,938,139 | 69,177,182 |
| *H. curtus* | chr7 | 30,195,766 | 22,850,312 | *H. elegans* | chr7 | 30,861,697 | 25,297,776 | *H. major* | chr7 | 42,362,139 | 20,915,223 |
| *H. curtus* | chr8 | 25,616,936 | 19,704,750 | *H. elegans* | chr8 | 25,217,042 | 20,632,258 | *H. major* | chr8 | 34,083,018 | 17,025,362 |
| *H. curtus* | chr9 | 25,286,039 | 18,978,143 | *H. elegans* | chr9 | 30,079,015 | 25,209,493 | *H. major* | chr9 | 30,848,814 | 20,497,424 |
| *H. curtus* | chr10 | 24,430,297 | 14,993,287 | *H. elegans* | chr10 | 23,601,959 | 19,245,899 | *H. major* | chr10 | 28,798,323 | 15,770,337 |
| *H. curtus* | chr11 | 22,723,820 | 17,432,170 | *H. elegans* | chr11 | 26,597,827 | 21,029,941 | *H. major* | chr11 | 28,504,944 | 17,354,881 |
| *H. curtus* | chr12 | 22,329,172 | 17,056,887 | *H. elegans* | chr12 | 16,702,216 | 12,699,160 | *H. major* | chr12 | 23,795,472 | 10,107,228 |
| *H. curtus* | chr13 | 19,757,143 | 14,406,772 | *H. elegans* | chr13 | 19,555,937 | 16,144,125 | *H. major* | chr13 | 19,560,250 | 13,307,042 |
| *H. curtus* | chr14 | 11,916,412 | 7,046,448 | *H. elegans* | chr14 | 11,861,178 | 8,816,035 | *H. major* | chr14 | 16,934,482 | 6,732,130 |
| *H. curtus* | chr15 | 7,845,892 | 4,616,978 | *H. elegans* | chr15 | 11,548,905 | 6,736,146 | *H. major* | chr15 | 16,659,334 | 5,517,956 |
| *H. curtus* | chr16 | 4,331,440 | 374,741 |  |  |  |  |  |  |  |  |
| Total | - | 1,733,272,931 | 1,284,653,362 | Total | - | 1,750,839,933 | 1,447,905,973 |  | Total | 1,793,465,688 | 1,189,863,120 |
| **Notes**<br>* All callable sites in this table are generated from reference-matched alignments that have been through the alignment and variant (and invariant) quality filtering pipelines. |  |  |  |  |  |  |  |  |  |  |  |

**Supplementary Table 5:**
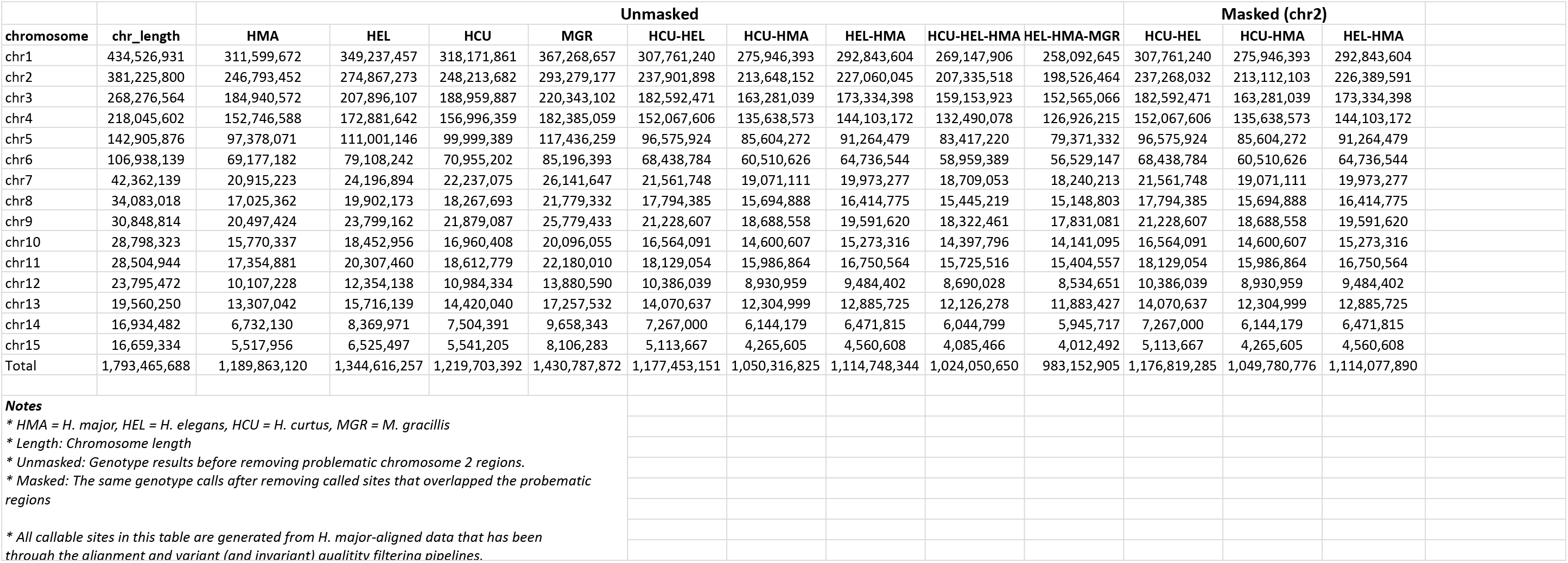
Callable sites along the *H. major* reference genome partitioned by chromosome.

|  |  | Unmasked |  |  |  |  |  |  |  |  | Masked (chr2) |  |  |
| --- | --- | --- | --- | --- | --- | --- | --- | --- | --- | --- | --- | --- | --- |
| chromosome | chr_length | HMA | HEL | HCU | MGR | HCU-HEL | HCU-HMA | HEL-HMA | HCU-HEL-HMA | HEL-HMA-MGR | HCU-HEL | HCU-HMA | HEL-HMA |
| chr1 | 434,526,931 | 311,599,672 | 349,237,457 | 318,171,861 | 367,268,657 | 307,761,240 | 275,946,393 | 292,843,604 | 269,147,906 | 258,092,645 | 307,761,240 | 275,946,393 | 292,843,604 |
| chr2 | 381,225,800 | 246,793,452 | 274,867,273 | 248,213,682 | 293,279,177 | 237,901,898 | 213,648,152 | 227,060,045 | 207,335,518 | 198,526,464 | 237,268,032 | 213,112,103 | 226,389,591 |
| chr3 | 268,276,564 | 184,940,572 | 207,896,107 | 188,959,887 | 220,343,102 | 182,592,471 | 163,281,039 | 173,334,398 | 159,153,923 | 152,565,066 | 182,592,471 | 163,281,039 | 173,334,398 |
| chr4 | 218,045,602 | 152,746,588 | 172,881,642 | 156,996,359 | 182,385,059 | 152,067,606 | 135,638,573 | 144,103,172 | 132,490,078 | 126,926,215 | 152,067,606 | 135,638,573 | 144,103,172 |
| chr5 | 142,905,876 | 97,378,071 | 111,001,146 | 99,999,389 | 117,436,259 | 96,575,924 | 85,604,272 | 91,264,479 | 83,417,220 | 79,371,332 | 96,575,924 | 85,604,272 | 91,264,479 |
| chr6 | 106,938,139 | 69,177,182 | 79,108,242 | 70,955,202 | 85,196,393 | 68,438,784 | 60,510,626 | 64,736,544 | 58,959,389 | 56,529,147 | 68,438,784 | 60,510,626 | 64,736,544 |
| chr7 | 42,362,139 | 20,915,223 | 24,196,894 | 22,237,075 | 26,141,647 | 21,561,748 | 19,071,111 | 19,973,277 | 18,709,053 | 18,240,213 | 21,561,748 | 19,071,111 | 19,973,277 |
| chr8 | 34,083,018 | 17,025,362 | 19,902,173 | 18,267,693 | 21,779,332 | 17,794,385 | 15,694,888 | 16,414,775 | 15,445,219 | 15,148,803 | 17,794,385 | 15,694,888 | 16,414,775 |
| chr9 | 30,848,814 | 20,497,424 | 23,799,162 | 21,879,087 | 25,779,433 | 21,228,607 | 18,688,558 | 19,591,620 | 18,322,461 | 17,831,081 | 21,228,607 | 18,688,558 | 19,591,620 |
| chr10 | 28,798,323 | 15,770,337 | 18,452,956 | 16,960,408 | 20,096,055 | 16,564,091 | 14,600,607 | 15,273,316 | 14,397,796 | 14,141,095 | 16,564,091 | 14,600,607 | 15,273,316 |
| chr11 | 28,504,944 | 17,354,881 | 20,307,460 | 18,612,779 | 22,180,010 | 18,129,054 | 15,986,864 | 16,750,564 | 15,725,516 | 15,404,557 | 18,129,054 | 15,986,864 | 16,750,564 |
| chr12 | 23,795,472 | 10,107,228 | 12,354,138 | 10,984,334 | 13,880,590 | 10,386,039 | 8,930,959 | 9,484,402 | 8,690,028 | 8,534,651 | 10,386,039 | 8,930,959 | 9,484,402 |
| chr13 | 19,560,250 | 13,307,042 | 15,716,139 | 14,420,040 | 17,257,532 | 14,070,637 | 12,304,999 | 12,885,725 | 12,126,278 | 11,883,427 | 14,070,637 | 12,304,999 | 12,885,725 |
| chr14 | 16,934,482 | 6,732,130 | 8,369,971 | 7,504,391 | 9,658,343 | 7,267,000 | 6,144,179 | 6,471,815 | 6,044,799 | 5,945,717 | 7,267,000 | 6,144,179 | 6,471,815 |
| chr15 | 16,659,334 | 5,517,956 | 6,525,497 | 5,541,205 | 8,106,283 | 5,113,667 | 4,265,605 | 4,560,608 | 4,085,466 | 4,012,492 | 5,113,667 | 4,265,605 | 4,560,608 |
| Total | 1,793,465,688 | 1,189,863,120 | 1,344,616,257 | 1,219,703,392 | 1,430,787,872 | 1,177,453,151 | 1,050,316,825 | 1,114,748,344 | 1,024,050,650 | 983,152,905 | 1,176,819,285 | 1,049,780,776 | 1,114,077,890 |
| Notes |  |  |  |  |  |  |  |  |  |  |  |  |  |
| * HMA = H. major, HEL = H. elegans, HCU = H. curtus, MGR = M. gracillis |  |  |  |  |  |  |  |  |  |  |  |  |  |
| * Length: Chromosome length |  |  |  |  |  |  |  |  |  |  |  |  |  |
| * Unmasked: Genotype results before removing problematic chromosome 2 regions. |  |  |  |  |  |  |  |  |  |  |  |  |  |
| * Masked: The same genotype calls after removing called sites that overlapped the problematic regions |  |  |  |  |  |  |  |  |  |  |  |  |  |
| * All callable sites in this table are generated from H. major-aligned data that has been through the alignment and variant (and invariant) quality filtering pipelines. |  |  |  |  |  |  |  |  |  |  |  |  |  |

**Supplementary table 6:**
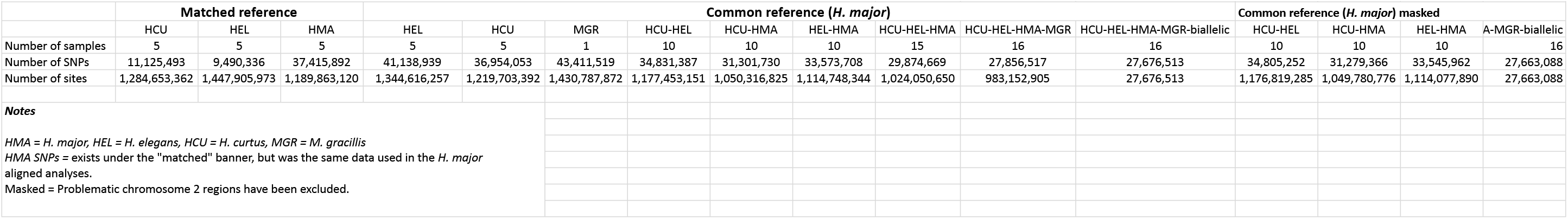
Called SNPs in matched and common reference datasets.

**Supplementary table 7:** Genome-scan outlier loci.

| Supplementary table 7: Genome-scan outlier loci |  |  |  |  |  |  |  |  |  |  |  |  |
| --- | --- | --- | --- | --- | --- | --- | --- | --- | --- | --- | --- | --- |
| comparison | chromosome | window_pos_1 | window_pos_2 | avg_wc_fst | no_snps | avg_dxy | no_sites | count_diffs | count_comparisons | count_missing | zfst | classification |
| HCU-HEL | 1 | 211,800,001 | 211,900,000 | 0.94 | 308 | 0.0020 | 73,528 | 13,876 | 6,843,696 | 509,104 | 3.07 | High |
| HCU-HEL | 1 | 212,400,001 | 212,500,000 | 0.94 | 470 | 0.0033 | 70,627 | 21,782 | 6,598,500 | 464,200 | 3.07 | High |
| HCU-HEL | 1 | 212,800,001 | 212,900,000 | 0.94 | 365 | 0.0031 | 56,140 | 16,266 | 5,230,476 | 383,524 | 3.07 | High |
| HCU-HEL | 1 | 213,600,001 | 213,700,000 | 0.94 | 492 | 0.0036 | 72,354 | 24,126 | 6,729,404 | 505,996 | 3.07 | High |
| HCU-HEL | 1 | 214,500,001 | 214,600,000 | 0.95 | 445 | 0.0029 | 76,811 | 20,926 | 7,115,100 | 566,000 | 3.21 | High |
| HCU-HEL | 1 | 214,600,001 | 214,700,000 | 0.97 | 445 | 0.0035 | 73,449 | 24,496 | 6,920,932 | 423,968 | 3.34 | High |
| HCU-HEL | 1 | 214,700,001 | 214,800,000 | 0.95 | 542 | 0.0041 | 74,928 | 28,922 | 6,985,472 | 507,328 | 3.18 | High |
| HCU-HEL | 1 | 214,800,001 | 214,900,000 | 0.95 | 503 | 0.0031 | 88,123 | 25,482 | 8,285,612 | 526,688 | 3.15 | High |
| HCU-HEL | 1 | 214,900,001 | 215,000,000 | 0.93 | 450 | 0.0033 | 66,357 | 20,542 | 6,248,356 | 387,344 | 3.04 | High |
| HCU-HEL | 2 | 238,500,001 | 238,600,000 | 0.93 | 434 | 0.0033 | 66,192 | 20,398 | 6,165,696 | 453,504 | 3.03 | High |
| HCU-HEL | 2 | 239,500,001 | 239,600,000 | 0.93 | 498 | 0.0043 | 65,516 | 25,866 | 6,055,900 | 495,700 | 3.02 | High |
| HCU-HEL | 2 | 240,800,001 | 240,900,000 | 0.94 | 374 | 0.0036 | 60,186 | 20,442 | 5,635,208 | 383,392 | 3.10 | High |
| HCU-HEL | 2 | 241,400,001 | 241,500,000 | 0.93 | 412 | 0.0033 | 63,360 | 19,924 | 5,955,580 | 380,420 | 3.03 | High |
| HCU-HEL | 3 | 121,000,001 | 121,100,000 | 0.95 | 492 | 0.0040 | 70,288 | 26,490 | 6,598,300 | 430,500 | 3.22 | High |
| HCU-HEL | 3 | 121,100,001 | 121,200,000 | 0.94 | 446 | 0.0034 | 72,344 | 22,742 | 6,782,340 | 452,060 | 3.10 | High |
| HCU-HEL | 3 | 123,100,001 | 123,200,000 | 0.93 | 307 | 0.0023 | 74,400 | 16,406 | 6,993,224 | 446,776 | 3.02 | High |
| HCU-HEL | 4 | 116,100,001 | 116,200,000 | 0.93 | 577 | 0.0042 | 72,704 | 28,328 | 6,792,196 | 478,204 | 3.00 | High |
| HCU-HEL | 4 | 116,600,001 | 116,700,000 | 0.94 | 743 | 0.0052 | 81,635 | 40,026 | 7,675,408 | 488,092 | 3.07 | High |
| HCU-HEL | 4 | 116,800,001 | 116,900,000 | 0.94 | 539 | 0.0037 | 83,437 | 29,176 | 7,916,688 | 427,012 | 3.14 | High |
| HCU-HEL | 4 | 156,400,001 | 156,500,000 | 0.21 | 1,403 | 0.0032 | 70,535 | 21,088 | 6,535,916 | 517,584 | -3.15 | Low |
| HCU-HEL | 5 | 64,700,001 | 64,800,000 | 0.93 | 338 | 0.0025 | 66,542 | 15,700 | 6,174,668 | 479,532 | 3.04 | High |
| HCU-HEL | 5 | 69,600,001 | 69,700,000 | 0.94 | 443 | 0.0036 | 70,245 | 23,516 | 6,571,960 | 452,540 | 3.14 | High |
| HCU-HEL | 5 | 71,000,001 | 71,100,000 | 0.93 | 407 | 0.0025 | 80,072 | 18,478 | 7,506,496 | 500,704 | 3.01 | High |
| HCU-HEL | 5 | 71,100,001 | 71,200,000 | 0.94 | 416 | 0.0027 | 77,446 | 19,628 | 7,252,800 | 491,800 | 3.09 | High |
| HCU-HEL | 5 | 72,200,001 | 72,300,000 | 0.93 | 466 | 0.0037 | 63,584 | 21,502 | 5,859,404 | 498,996 | 3.01 | High |
| HCU-HEL | 6 | 5,800,001 | 5,900,000 | 0.22 | 1,363 | 0.0063 | 44,274 | 25,456 | 4,012,256 | 415,144 | -3.03 | Low |
| HCU-HEL | 6 | 5,900,001 | 6,000,000 | 0.18 | 1,433 | 0.0057 | 48,499 | 24,804 | 4,372,364 | 477,536 | -3.41 | Low |
| HCU-HEL | 6 | 6,100,001 | 6,200,000 | 0.21 | 2,138 | 0.0064 | 67,158 | 38,944 | 6,124,068 | 591,732 | -3.15 | Low |
| HCU-HEL | 6 | 6,300,001 | 6,400,000 | 0.21 | 344 | 0.0060 | 11,866 | 5,560 | 930,540 | 256,060 | -3.14 | Low |
| HCU-HEL | 6 | 92,500,001 | 92,600,000 | 0.96 | 486 | 0.0038 | 66,484 | 22,782 | 5,984,752 | 663,648 | 3.27 | High |
| HCU-HEL | 6 | 92,700,001 | 92,800,000 | 0.95 | 282 | 0.0036 | 42,662 | 13,968 | 3,930,820 | 335,380 | 3.20 | High |
| HCU-HEL | 6 | 92,900,001 | 93,000,000 | 0.95 | 395 | 0.0030 | 72,058 | 20,298 | 6,783,384 | 422,416 | 3.22 | High |
| HCU-HEL | 6 | 94,000,001 | 94,100,000 | 0.94 | 124 | 0.0062 | 12,915 | 7,318 | 1,181,572 | 109,928 | 3.06 | High |
| HCU-HEL | 6 | 94,300,001 | 94,400,000 | 0.97 | 383 | 0.0037 | 66,189 | 22,856 | 6,191,896 | 427,004 | 3.35 | High |
| HCU-HEL | 9 | 1 | 100,000 | 0.94 | 264 | 0.0051 | 31,977 | 15,326 | 2,986,848 | 210,852 | 3.10 | High |
| HCU-HEL | 13 | 200,001 | 300,000 | 0.93 | 170 | 0.0035 | 23,127 | 7,714 | 2,182,060 | 130,640 | 3.05 | High |
| HCU-HEL | 13 | 300,001 | 400,000 | 0.95 | 472 | 0.0034 | 74,927 | 24,040 | 7,060,788 | 431,912 | 3.23 | High |
| HCU-HMA | 1 | 211,800,001 | 211,900,000 | 0.94 | 213 | 0.0024 | 64,793 | 13,856 | 5,692,752 | 786,548 | 3.06 | High |
| HCU-HMA | 1 | 212,300,001 | 212,400,000 | 0.93 | 319 | 0.0034 | 68,721 | 20,626 | 6,099,056 | 773,044 | 3.00 | High |
| HCU-HMA | 1 | 212,400,001 | 212,500,000 | 0.95 | 360 | 0.0042 | 65,953 | 24,526 | 5,847,728 | 747,572 | 3.17 | High |
| HCU-HMA | 1 | 212,600,001 | 212,700,000 | 0.93 | 407 | 0.0043 | 68,709 | 26,152 | 6,148,384 | 722,516 | 3.01 | High |
| HCU-HMA | 1 | 212,700,001 | 212,800,000 | 0.95 | 362 | 0.0054 | 52,831 | 25,266 | 4,662,504 | 620,596 | 3.18 | High |
| HCU-HMA | 1 | 212,800,001 | 212,900,000 | 0.96 | 248 | 0.0038 | 51,751 | 17,134 | 4,553,388 | 621,712 | 3.26 | High |
| HCU-HMA | 1 | 213,000,001 | 213,100,000 | 0.93 | 348 | 0.0037 | 69,478 | 22,410 | 6,125,688 | 822,112 | 3.03 | High |
| HCU-HMA | 1 | 213,600,001 | 213,700,000 | 0.94 | 374 | 0.0040 | 66,751 | 24,114 | 5,965,260 | 709,840 | 3.06 | High |
| HCU-HMA | 1 | 214,100,001 | 214,200,000 | 0.94 | 310 | 0.0038 | 60,082 | 19,978 | 5,262,672 | 745,528 | 3.11 | High |
| HCU-HMA | 1 | 214,300,001 | 214,400,000 | 0.96 | 272 | 0.0032 | 68,724 | 19,548 | 6,034,780 | 837,620 | 3.29 | High |
| HCU-HMA | 1 | 214,500,001 | 214,600,000 | 0.96 | 261 | 0.0031 | 68,282 | 18,820 | 6,001,424 | 826,776 | 3.33 | High |
| HCU-HMA | 1 | 214,600,001 | 214,700,000 | 0.97 | 246 | 0.0032 | 65,913 | 18,674 | 5,897,276 | 694,024 | 3.40 | High |
| HCU-HMA | 1 | 214,700,001 | 214,800,000 | 0.95 | 319 | 0.0036 | 69,383 | 22,112 | 6,166,888 | 771,412 | 3.18 | High |
| HCU-HMA | 1 | 214,800,001 | 214,900,000 | 0.94 | 312 | 0.0029 | 80,975 | 20,722 | 7,212,176 | 885,324 | 3.08 | High |
| HCU-HMA | 1 | 228,200,001 | 228,300,000 | 0.29 | 919 | 0.0055 | 44,799 | 20,572 | 3,745,172 | 734,728 | -3.42 | Low |
| HCU-HMA | 1 | 398,900,001 | 399,000,000 | 0.94 | 646 | 0.0092 | 55,425 | 45,674 | 4,954,644 | 587,856 | 3.08 | High |
| HCU-HMA | 2 | 31,500,001 | 31,600,000 | 0.33 | 855 | 0.0053 | 45,354 | 21,018 | 3,996,404 | 538,996 | -3.04 | Low |
| HCU-HMA | 3 | 124,100,001 | 124,200,000 | 0.93 | 246 | 0.0030 | 59,461 | 15,776 | 5,337,084 | 609,016 | 3.02 | High |
| HCU-HMA | 4 | 4,300,001 | 4,400,000 | 0.31 | 1,620 | 0.0070 | 62,812 | 39,082 | 5,546,312 | 734,888 | -3.23 | Low |
| HCU-HMA | 4 | 116,800,001 | 116,900,000 | 0.95 | 382 | 0.0037 | 75,807 | 25,158 | 6,804,196 | 776,504 | 3.15 | High |
| HCU-HMA | 5 | 66,000,001 | 66,100,000 | 0.93 | 259 | 0.0030 | 65,228 | 17,050 | 5,756,208 | 766,592 | 3.02 | High |
| HCU-HMA | 5 | 68,700,001 | 68,800,000 | 0.94 | 407 | 0.0053 | 57,967 | 26,836 | 5,082,992 | 713,708 | 3.13 | High |
| HCU-HMA | 5 | 69,200,001 | 69,300,000 | 0.93 | 231 | 0.0039 | 43,625 | 14,498 | 3,752,456 | 610,044 | 3.02 | High |
| HCU-HMA | 5 | 69,600,001 | 69,700,000 | 0.94 | 280 | 0.0037 | 59,246 | 19,344 | 5,287,252 | 637,348 | 3.10 | High |
| HCU-HMA | 6 | 5,900,001 | 6,000,000 | 0.31 | 981 | 0.0052 | 40,662 | 17,896 | 3,431,456 | 634,744 | -3.23 | Low |
| HCU-HMA | 6 | 6,000,001 | 6,100,000 | 0.27 | 1,124 | 0.0056 | 44,949 | 21,766 | 3,895,048 | 599,852 | -3.56 | Low |
| HCU-HMA | 6 | 6,100,001 | 6,200,000 | 0.33 | 1,491 | 0.0062 | 59,580 | 32,014 | 5,158,860 | 799,140 | -3.02 | Low |
| HCU-HMA | 6 | 6,200,001 | 6,300,000 | 0.30 | 912 | 0.0054 | 40,193 | 19,382 | 3,570,100 | 449,200 | -3.26 | Low |
| HCU-HMA | 6 | 6,300,001 | 6,400,000 | 0.29 | 206 | 0.0045 | 10,185 | 3,484 | 773,904 | 244,596 | -3.40 | Low |
| HCU-HMA | 6 | 6,600,001 | 6,700,000 | 0.30 | 1,553 | 0.0077 | 57,704 | 38,764 | 5,032,344 | 738,056 | -3.33 | Low |
| HCU-HMA | 6 | 92,500,001 | 92,600,000 | 0.96 | 277 | 0.0045 | 51,837 | 19,830 | 4,383,952 | 799,748 | 3.29 | High |
| HCU-HMA | 6 | 92,700,001 | 92,800,000 | 0.96 | 165 | 0.0038 | 36,337 | 11,968 | 3,188,640 | 445,060 | 3.31 | High |
| HCU-HMA | 6 | 92,800,001 | 92,900,000 | 0.94 | 436 | 0.0046 | 72,328 | 29,484 | 6,373,504 | 859,296 | 3.08 | High |
| HCU-HMA | 6 | 92,900,001 | 93,000,000 | 0.96 | 257 | 0.0035 | 62,387 | 19,352 | 5,604,904 | 633,796 | 3.32 | High |
| HCU-HMA | 6 | 93,800,001 | 93,900,000 | 0.95 | 217 | 0.0079 | 22,363 | 14,914 | 1,881,600 | 354,700 | 3.22 | High |
| HCU-HMA | 6 | 94,000,001 | 94,100,000 | 0.94 | 129 | 0.0073 | 13,840 | 8,882 | 1,215,264 | 168,736 | 3.05 | High |
| HCU-HMA | 6 | 94,100,001 | 94,200,000 | 0.94 | 131 | 0.0073 | 14,646 | 9,384 | 1,283,796 | 180,804 | 3.07 | High |
| HCU-HMA | 6 | 94,300,001 | 94,400,000 | 0.96 | 279 | 0.0041 | 59,016 | 21,428 | 5,268,472 | 633,128 | 3.33 | High |
| HCU-HMA | 9 | 1 | 100,000 | 0.93 | 199 | 0.0055 | 26,974 | 13,248 | 2,403,432 | 293,968 | 3.01 | High |
| HCU-HMA | 11 | 28,200,001 | 28,300,000 | 0.94 | 443 | 0.0049 | 68,751 | 29,360 | 6,002,252 | 872,848 | 3.08 | High |
| HCU-HMA | 11 | 28,300,001 | 28,400,000 | 0.93 | 353 | 0.0037 | 69,546 | 22,336 | 6,051,392 | 903,208 | 3.02 | High |
| HCU-HMA | 13 | 200,001 | 300,000 | 0.93 | 132 | 0.0044 | 21,437 | 8,500 | 1,932,720 | 210,980 | 3.02 | High |
| HCU-HMA | 13 | 300,001 | 400,000 | 0.96 | 329 | 0.0038 | 68,448 | 23,392 | 6,173,660 | 671,140 | 3.25 | High |
| HEL-HMA | 1 | 9,400,001 | 9,500,000 | 0.36 | 1,274 | 0.0054 | 65,768 | 32,492 | 5,992,048 | 584,752 | -3.39 | Low |
| HEL-HMA | 1 | 134,700,001 | 134,800,000 | 0.13 | 253 | 0.0069 | 10,870 | 6,006 | 874,952 | 212,048 | -5.84 | Low |
| HEL-HMA | 1 | 135,000,001 | 135,100,000 | 0.29 | 457 | 0.0053 | 27,519 | 12,956 | 2,428,992 | 322,908 | -4.15 | Low |
| HEL-HMA | 1 | 214,500,001 | 214,600,000 | 0.98 | 327 | 0.0039 | 72,453 | 25,384 | 6,537,188 | 708,112 | 3.10 | High |
| HEL-HMA | 1 | 214,600,001 | 214,700,000 | 0.98 | 331 | 0.0041 | 70,012 | 26,258 | 6,397,576 | 603,624 | 3.13 | High |
| HEL-HMA | 2 | 20,200,001 | 20,300,000 | 0.36 | 609 | 0.0036 | 72,324 | 23,886 | 6,564,008 | 668,392 | -3.42 | Low |
| HEL-HMA | 2 | 20,300,001 | 20,400,000 | 0.36 | 417 | 0.0030 | 54,970 | 14,708 | 4,945,340 | 551,660 | -3.45 | Low |
| HEL-HMA | 2 | 20,600,001 | 20,700,000 | 0.26 | 154 | 0.0039 | 10,801 | 3,392 | 869,164 | 210,936 | -4.41 | Low |
| HEL-HMA | 2 | 21,600,001 | 21,700,000 | 0.32 | 199 | 0.0042 | 15,007 | 5,154 | 1,226,160 | 274,540 | -3.81 | Low |
| HEL-HMA | 2 | 23,000,001 | 23,100,000 | 0.37 | 420 | 0.0038 | 40,521 | 13,714 | 3,606,504 | 445,596 | -3.34 | Low |
| HEL-HMA | 2 | 25,900,001 | 26,000,000 | 0.40 | 324 | 0.0045 | 25,448 | 9,920 | 2,183,144 | 361,656 | -3.00 | Low |
| HEL-HMA | 2 | 26,100,001 | 26,200,000 | 0.19 | 365 | 0.0041 | 28,616 | 9,288 | 2,242,504 | 619,096 | -5.14 | Low |
| HEL-HMA | 2 | 26,300,001 | 26,400,000 | 0.36 | 336 | 0.0063 | 23,467 | 11,678 | 1,843,252 | 503,448 | -3.37 | Low |
| HEL-HMA | 2 | 26,400,001 | 26,500,000 | 0.29 | 669 | 0.0032 | 68,783 | 19,790 | 6,212,784 | 665,516 | -4.15 | Low |
| HEL-HMA | 2 | 30,200,001 | 30,300,000 | 0.28 | 357 | 0.0073 | 14,882 | 8,684 | 1,188,172 | 300,028 | -4.19 | Low |
| HEL-HMA | 2 | 31,500,001 | 31,600,000 | 0.35 | 809 | 0.0053 | 48,026 | 23,032 | 4,343,544 | 459,056 | -3.48 | Low |
| HEL-HMA | 2 | 31,800,001 | 31,900,000 | 0.31 | 1,040 | 0.0067 | 46,712 | 27,212 | 4,072,308 | 598,892 | -3.93 | Low |
| HEL-HMA | 2 | 37,000,001 | 37,100,000 | 0.40 | 207 | 0.0061 | 10,503 | 5,114 | 841,852 | 208,448 | -3.02 | Low |
| HEL-HMA | 2 | 39,100,001 | 39,200,000 | 0.30 | 360 | 0.0049 | 18,572 | 7,452 | 1,513,288 | 343,912 | -4.07 | Low |
| HEL-HMA | 2 | 51,500,001 | 51,600,000 | 0.35 | 172 | 0.0034 | 13,845 | 3,450 | 1,022,936 | 361,564 | -3.49 | Low |
| HEL-HMA | 2 | 51,600,001 | 51,700,000 | 0.39 | 183 | 0.0037 | 14,916 | 4,196 | 1,147,008 | 344,592 | -3.09 | Low |
| HEL-HMA | 2 | 52,100,001 | 52,200,000 | 0.29 | 934 | 0.0059 | 52,906 | 27,592 | 4,650,416 | 640,184 | -4.10 | Low |
| HEL-HMA | 2 | 52,200,001 | 52,300,000 | 0.23 | 601 | 0.0052 | 33,572 | 15,140 | 2,886,604 | 470,596 | -4.73 | Low |
| HEL-HMA | 2 | 52,300,001 | 52,400,000 | 0.34 | 1,193 | 0.0061 | 61,997 | 34,474 | 5,634,652 | 565,048 | -3.64 | Low |
| HEL-HMA | 2 | 59,700,001 | 59,800,000 | 0.37 | 520 | 0.0051 | 30,679 | 13,086 | 2,550,436 | 517,464 | -3.26 | Low |
| HEL-HMA | 2 | 61,900,001 | 62,000,000 | 0.38 | 772 | 0.0049 | 40,841 | 17,950 | 3,642,788 | 441,312 | -3.21 | Low |
| HEL-HMA | 2 | 225,200,001 | 225,300,000 | 0.39 | 710 | 0.0104 | 26,011 | 23,554 | 2,259,272 | 341,828 | -3.13 | Low |
| HEL-HMA | 2 | 227,400,001 | 227,500,000 | 0.40 | 1,314 | 0.0047 | 75,666 | 32,110 | 6,880,020 | 686,580 | -3.02 | Low |
| HEL-HMA | 2 | 349,800,001 | 349,900,000 | 0.34 | 785 | 0.0042 | 58,323 | 22,658 | 5,357,288 | 475,012 | -3.59 | Low |
| HEL-HMA | 3 | 253,300,001 | 253,400,000 | 0.38 | 1,321 | 0.0059 | 69,150 | 37,130 | 6,340,248 | 574,752 | -3.17 | Low |
| HEL-HMA | 3 | 263,500,001 | 263,600,000 | 0.38 | 1,326 | 0.0066 | 60,760 | 35,858 | 5,435,384 | 640,616 | -3.22 | Low |
| HEL-HMA | 3 | 267,900,001 | 268,000,000 | 0.39 | 1,048 | 0.0077 | 41,336 | 28,604 | 3,717,500 | 416,100 | -3.12 | Low |
| HEL-HMA | 4 | 3,200,001 | 3,300,000 | 0.39 | 1,365 | 0.0073 | 57,814 | 37,920 | 5,204,540 | 576,860 | -3.05 | Low |
| HEL-HMA | 4 | 4,300,001 | 4,400,000 | 0.38 | 1,637 | 0.0075 | 67,407 | 45,256 | 6,051,484 | 689,216 | -3.13 | Low |
| HEL-HMA | 4 | 29,100,001 | 29,200,000 | 0.39 | 991 | 0.0035 | 74,603 | 24,012 | 6,803,340 | 656,960 | -3.05 | Low |
| HEL-HMA | 4 | 87,300,001 | 87,400,000 | 0.34 | 1,178 | 0.0061 | 57,814 | 31,404 | 5,136,356 | 645,044 | -3.58 | Low |
| HEL-HMA | 4 | 212,900,001 | 213,000,000 | 0.34 | 1,390 | 0.0079 | 50,118 | 35,662 | 4,518,560 | 493,240 | -3.58 | Low |
| HEL-HMA | 6 | 6,000,001 | 6,100,000 | 0.37 | 1,033 | 0.0058 | 49,321 | 25,452 | 4,397,236 | 534,864 | -3.25 | Low |
| HEL-HMA | 6 | 6,100,001 | 6,200,000 | 0.37 | 1,526 | 0.0063 | 66,594 | 37,420 | 5,987,060 | 672,340 | -3.32 | Low |
| HEL-HMA | 6 | 84,900,001 | 85,000,000 | 0.31 | 256 | 0.0040 | 17,481 | 5,886 | 1,482,744 | 265,356 | -3.93 | Low |
| HEL-HMA | 6 | 95,500,001 | 95,600,000 | 0.37 | 1,064 | 0.0051 | 65,570 | 30,284 | 5,971,296 | 585,704 | -3.32 | Low |
| HEL-HMA | 6 | 95,600,001 | 95,700,000 | 0.37 | 977 | 0.0043 | 73,737 | 29,130 | 6,704,888 | 668,812 | -3.27 | Low |
| HEL-HMA | 15 | 9,100,001 | 9,200,000 | 0.40 | 1,135 | 0.0078 | 43,909 | 30,636 | 3,909,540 | 481,360 | -3.02 | Low |

**Supplementary table 8:**
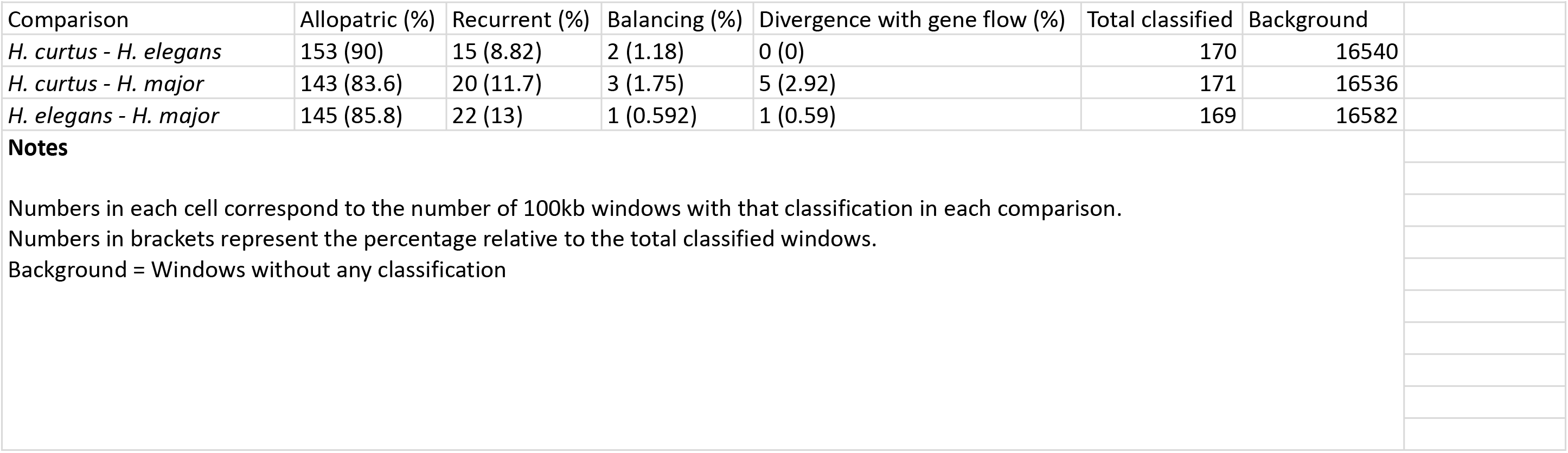
Divergence profiling summary.

**Supplementary table 9:**
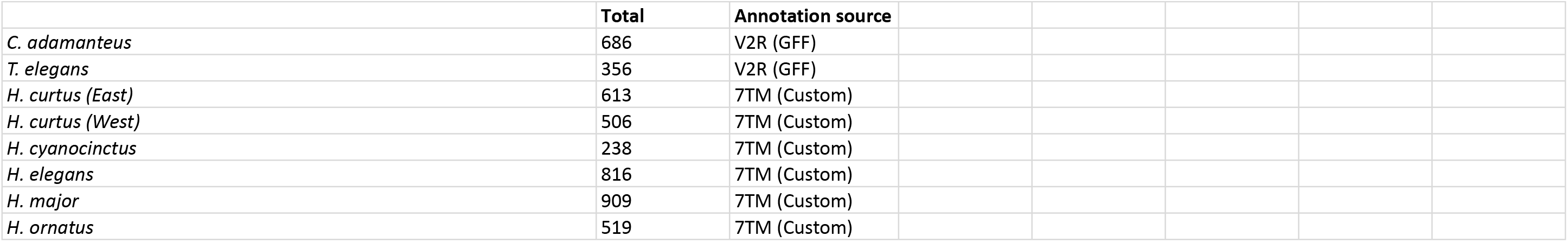
Total V2R (7TM) content in explored snakes.

**Supplementary table 10:** V2R (7TM) content by chromosome.

| Supplementary table 10: V2R (7TM) content by chromosome |  |  |  |  |  |  |  |  |
| --- | --- | --- | --- | --- | --- | --- | --- | --- |
|  | *H. curtus (East)* | *H. curtus (West)* | *H. cyanocinctus* | *H. elegans* | *H. major* | *H. ornatus* | *C. adamanteus* | *T. elegans* |
| chr1 | 31 | 78 | 20 | 66 | 68 | 56 | 48 | 52 |
| chr2 | 300 | 247 | 48 | 524 | 556 | 270 | 279 | 28 |
| chr3 | 1 | 1 | 1 | 1 | 1 | 11 |  |  |
| chr4 | 16 | 11 | 3 | 14 | 10 | 2 |  |  |
| chr5 |  |  |  |  |  |  | 1 |  |
| chr6 |  | 1 |  | 3 | 3 | 3 |  |  |
| chr7 |  | 1 |  | 3 | 1 |  |  |  |
| chr8 |  |  |  | 2 | 9 |  |  |  |
| chr9 |  |  |  |  |  | 1 |  |  |
| chr10 |  |  |  | 6 | 2 |  |  | 1 |
| chr11 | 26 |  |  |  |  |  |  |  |
| chr12 |  |  |  | 5 | 32 |  |  |  |
| chr14 |  |  | 1 | 1 |  |  |  |  |
| chr15 | 1 | 1 | 4 | 21 | 55 | 8 | 7 |  |
| chr16 |  |  | 1 |  |  |  | 12 |  |
| chr17 | 31 |  |  |  |  |  |  |  |
| chr18 |  |  | 21 |  |  |  |  |  |
| chrZ | 207 | 166 | 139 | 170 | 172 | 168 | 338 | 242 |
| chrW |  |  |  |  |  |  | 1 |  |
| chrUn |  |  |  |  |  |  |  | 33 |

## Notes

### Competing Interest Statement

The authors have declared no competing interest.

## References

Alonge, M., Lebeigle, L., Kirsche, M., Jenike, K., Ou, S., Aganezov, S., Wang, X., Lippman, Z. B., Schatz, M. C., & Soyk, S. (2022). Automated assembly scaffolding using RagTag elevates a new tomato system for high-throughput genome editing. Genome Biology, 23(1), 258. 10.1186/s13059-022-02823-7

Alonso, S., López, S., Izagirre, N., & de la Rúa, C. (2008). Overdominance in the Human Genome and Olfactory Receptor Activity. Molecular Biology and Evolution, 25(5), 997–1001. 10.1093/molbev/msn049

Andrew, R. L., & Rieseberg, L. H. (2013). Divergence Is Focused on Few Genomic Regions Early in Speciation: Incipient Speciation of Sunflower Ecotypes. Evolution, 67(9), 2468–2482. 10.1111/evo.12106

Brykczynska, U., Tzika, A. C., Rodriguez, I., & Milinkovitch, M. C. (2013). Contrasted Evolution of the Vomeronasal Receptor Repertoires in Mammals and Squamate Reptiles. Genome Biology and Evolution, 5(2), 389–401. 10.1093/gbe/evt013

Burri, R. (2017). Interpreting differentiation landscapes in the light of long-term linked selection. Evolution Letters, 1(3), 118–131. 10.1002/evl3.14

C. Fry, G., A. Milton, D., & J. Wassenberg, T. (2001). The reproductive biology and diet of sea snake bycatch of prawn trawling in northern Australia: Characteristics important for assessing the impacts on populations. Pacific Conservation Biology, 7(1), 55–73. 10.1071/PC010055

Camacho, C., Coulouris, G., Avagyan, V., Ma, N., Papadopoulos, J., Bealer, K., & Madden, T. L. (2009). BLAST+: Architecture and applications. BMC Bioinformatics, 10, 421. 10.1186/1471-2105-10-421

Chen, S. (2023). Ultrafast one-pass FASTQ data preprocessing, quality control, and deduplication using fastp. iMeta, 2(2), e107. 10.1002/imt2.107

Cheng, S., & Faidi, M. A. (2025). Palaeodrainages of the Sunda Shelf detailed in new maps. Journal of Palaeogeography, 14(1), 186–202. 10.1016/j.jop.2024.10.001

Christmas, M. J., Jones, J. C., Olsson, A., Wallerman, O., Bunikis, I., Kierczak, M., Peona, V., Whitley, K. M., Larva, T., Suh, A., Miller-Struttmann, N. E., Geib, J. C., & Webster, M. T. (2021). Genetic Barriers to Historical Gene Flow between Cryptic Species of Alpine Bumblebees Revealed by Comparative Population Genomics. Molecular Biology and Evolution, 38(8), 3126–3143. 10.1093/molbev/msab086

Croze, M., Živković, D., Stephan, W., & Hutter, S. (2016). Balancing selection on immunity genes: Review of the current literature and new analysis in *Drosophila melanogaster*. *Zoology*, SI: Host-Parasite Coevolution, 119(4), 322–329. 10.1016/j.zool.2016.03.004

Cruickshank, T. E., & Hahn, M. W. (2014). Reanalysis suggests that genomic islands of speciation are due to reduced diversity, not reduced gene flow. Molecular Ecology, 23(13), 3133–3157. 10.1111/mec.12796

Dainat, J. (2024). NBISweden/AGAT: AGAT-v1.4.0 (Version v1.4.0) [Computer software]. Zenodo. 10.5281/zenodo.11106497

Danecek, P., Auton, A., Abecasis, G., Albers, C. A., Banks, E., DePristo, M. A., Handsaker, R. E., Lunter, G., Marth, G. T., Sherry, S. T., McVean, G., Durbin, R., & 1000 Genomes Project Analysis Group. (2011). The variant call format and VCFtools. Bioinformatics, 27(15), 2156–2158. 10.1093/bioinformatics/btr330

Danecek, P., Bonfield, J. K., Liddle, J., Marshall, J., Ohan, V., Pollard, M. O., Whitwham, A., Keane, T., McCarthy, S. A., Davies, R. M., & Li, H. (2021). Twelve years of SAMtools and BCFtools. GigaScience, 10(2), giab008. 10.1093/gigascience/giab008

Eddy, S. R. (2011). Accelerated Profile HMM Searches. PLOS Computational Biology, 7(10), e1002195. 10.1371/journal.pcbi.1002195

Ellegren, H., Smeds, L., Burri, R., Olason, P. I., Backström, N., Kawakami, T., Künstner, A., Mäkinen, H., Nadachowska-Brzyska, K., Qvarnström, A., Uebbing, S., & Wolf, J. B. W. (2012). The genomic landscape of species divergence in Ficedula flycatchers. Nature, 491(7426), 756–760. 10.1038/nature11584

Feder, J. L., Egan, S. P., & Nosil, P. (2012). The genomics of speciation-with-gene-flow. Trends in Genetics, 28(7), 342–350. 10.1016/j.tig.2012.03.009

Fu, Y. X. (1995). Statistical Properties of Segregating Sites. Theoretical Population Biology, 48(2), 172–197. 10.1006/tpbi.1995.1025

Gel, B., Díez-Villanueva, A., Serra, E., Buschbeck, M., Peinado, M. A., & Malinverni, R. (2016). regioneR: An R/Bioconductor package for the association analysis of genomic regions based on permutation tests. Bioinformatics, 32(2), 289–291. 10.1093/bioinformatics/btv562

Han, F., Lamichhaney, S., Grant, B. R., Grant, P. R., Andersson, L., & Webster, M. T. (2017). Gene flow, ancient polymorphism, and ecological adaptation shape the genomic landscape of divergence among Darwin’s finches. Genome Research, 27(6), 1004–1015. 10.1101/gr.212522.116

Hedrick, P. W. (2007). Balancing selection. Current Biology, 17(7), R230–R231. 10.1016/j.cub.2007.01.012

Hogan, M. P., Holding, M. L., Nystrom, G. S., Lawrence, K. C., Broussard, E. M., Ellsworth, S. A., Mason, A. J., Margres, M. J., Gibbs, H. L., Parkinson, C. L., & Rokyta, D. R. (2025). Life history and chromosome organization determine chemoreceptor gene expression in rattlesnakes. Journal of Heredity, 116(5), 617–631. 10.1093/jhered/esae078

Hogan, M. P., Whittington, A. C., Broe, M. B., Ward, M. J., Gibbs, H. L., & Rokyta, D. R. (2021). The Chemosensory Repertoire of the Eastern Diamondback Rattlesnake (Crotalus adamanteus) Reveals Complementary Genetics of Olfactory and Vomeronasal-Type Receptors. Journal of Molecular Evolution, 89(4), 313–328. 10.1007/s00239-021-10007-3

Holsinger, K. E., & Weir, B. S. (2009). Genetics in geographically structured populations: Defining, estimating and interpreting FST. Nature Reviews Genetics, 10(9), 639–650. 10.1038/nrg2611

Irwin, D. E., Milá, B., Toews, D. P. L., Brelsford, A., Kenyon, H. L., Porter, A. N., Grossen, C., Delmore, K. E., Alcaide, M., & Irwin, J. H. (2018). A comparison of genomic islands of differentiation across three young avian species pairs. Molecular Ecology, 27(23), 4839–4855. 10.1111/mec.14858

Kautt, A. F., Kratochwil, C. F., Nater, A., Machado-Schiaffino, G., Olave, M., Henning, F., Torres-Dowdall, J., Härer, A., Hulsey, C. D., Franchini, P., Pippel, M., Myers, E. W., & Meyer, A. (2020). Contrasting signatures of genomic divergence during sympatric speciation. Nature, 588(7836), 106–111. 10.1038/s41586-020-2845-0

Kishida, T., Keboushi, R., Sasai, T., & Toda, M. (2025). Genomics Reveals Recent Rapid Speciation of Sea Snakes of the Genus Hydrophis (Reptilia, Squamata, Elapidae). Ecology and Evolution, 15(6), e71627. 10.1002/ece3.71627

Kishida, T., Toda, M., Go, Y., Tatsumoto, S., Sasai, T., & Hikida, T. (2021). Population history and genomic admixture of sea snakes of the genus *Laticauda* in the West Pacific. Molecular Phylogenetics and Evolution, 155, 107005. 10.1016/j.ympev.2020.107005

Korneliussen, T. S., Albrechtsen, A., & Nielsen, R. (2014). ANGSD: Analysis of Next Generation Sequencing Data. BMC Bioinformatics, 15(1), 356. 10.1186/s12859-014-0356-4

Korunes, K. L., & Samuk, K. (2021). pixy: Unbiased estimation of nucleotide diversity and divergence in the presence of missing data. Molecular Ecology Resources, 21(4), 1359–1368. 10.1111/1755-0998.13326

Kronforst, M. R., Hansen, M. E. B., Crawford, N. G., Gallant, J. R., Zhang, W., Kulathinal, R. J., Kapan, D. D., & Mullen, S. P. (2013). Hybridization Reveals the Evolving Genomic Architecture of Speciation. Cell Reports, 5(3), 666–677. 10.1016/j.celrep.2013.09.042

Kutsuma, R., Sasai, T., & Kishida, T. (2018). How Snakes Find Prey Underwater: Sea Snakes Use Visual and Chemical Cues for Foraging. Zoological Science, 35(6), 483–486. 10.2108/zs180059

Lee, M. S. Y., Sanders, K. L., King, B., & Palci, A. (2016). Diversification rates and phenotypic evolution in venomous snakes (Elapidae). Royal Society Open Science, 3(1), 150277. 10.1098/rsos.150277

Li, A., Wang, J., Sun, K., Wang, S., Zhao, X., Wang, T., Xiong, L., Xu, W., Qiu, L., Shang, Y., Liu, R., Wang, S., & Lu, Y. (2021). Two Reference-Quality Sea Snake Genomes Reveal Their Divergent Evolution of Adaptive Traits and Venom Systems. Molecular Biology and Evolution, 38(11), 4867–4883. 10.1093/molbev/msab212

Li, H. (2023). Protein-to-genome alignment with miniprot. Bioinformatics, 39(1), btad014. 10.1093/bioinformatics/btad014

Li, W., & Godzik, A. (2006). Cd-hit: A fast program for clustering and comparing large sets of protein or nucleotide sequences. Bioinformatics, 22(13), 1658–1659. 10.1093/bioinformatics/btl158

Lin, Y., Ye, C., Li, X., Chen, Q., Wu, Y., Zhang, F., Pan, R., Zhang, S., Chen, S., Wang, X., Cao, S., Wang, Y., Yue, Y., Liu, Y., & Yue, J. (2023). quarTeT: A telomere-to-telomere toolkit for gap-free genome assembly and centromeric repeat identification. Horticulture Research, 10(8), uhad127. 10.1093/hr/uhad127

Liu, X., Glémin, S., & Karrenberg, S. (2020). Evolution of putative barrier loci at an intermediate stage of speciation with gene flow in campions (Silene). Molecular Ecology, 29(18), 3511–3525. 10.1111/mec.15571

Ludington, A. J., Hammond, J. M., Breen, J., Deveson, I. W., & Sanders, K. L. (2023). New chromosome-scale genomes provide insights into marine adaptations of sea snakes (Hydrophis: Elapidae). BMC Biology, 21(1), 284. 10.1186/s12915-023-01772-2

Ludington, A. J., & Sanders, K. L. (2021). Demographic analyses of marine and terrestrial snakes (Elapidae) using whole genome sequences. Molecular Ecology, 30(2), 545–554. 10.1111/mec.15726

Ludt, W. B., & Rocha, L. A. (2015). Shifting seas: The impacts of Pleistocene sea-level fluctuations on the evolution of tropical marine taxa. Journal of Biogeography, 42(1), 25–38. 10.1111/jbi.12416

Ma, T., Wang, K., Hu, Q., Xi, Z., Wan, D., Wang, Q., Feng, J., Jiang, D., Ahani, H., Abbott, R. J., Lascoux, M., Nevo, E., & Liu, J. (2018). Ancient polymorphisms and divergence hitchhiking contribute to genomic islands of divergence within a poplar species complex. Proceedings of the National Academy of Sciences, 115(2). 10.1073/pnas.1713288114

Malinsky, M., Matschiner, M., & Svardal, H. (2021). Dsuite—Fast D-statistics and related admixture evidence from VCF files. Molecular Ecology Resources, 21(2), 584–595. 10.1111/1755-0998.13265

Malnic, B., Hirono, J., Sato, T., & Buck, L. B. (1999). Combinatorial Receptor Codes for Odors. Cell, 96(5), 713–723. 10.1016/S0092-8674(00)80581-4

Martin, S. H., Dasmahapatra, K. K., Nadeau, N. J., Salazar, C., Walters, J. R., Simpson, F., Blaxter, M., Manica, A., Mallet, J., & Jiggins, C. D. (2013). Genome-wide evidence for speciation with gene flow in Heliconius butterflies. Genome Research. 10.1101/gr.159426.113

Martin, S. H., Davey, J. W., & Jiggins, C. D. (2015). Evaluating the Use of ABBA–BABA Statistics to Locate Introgressed Loci. Molecular Biology and Evolution, 32(1), 244–257. 10.1093/molbev/msu269

Martin, S. H., Möst, M., Palmer, W. J., Salazar, C., McMillan, W. O., Jiggins, F. M., & Jiggins, C. D. (2016). Natural Selection and Genetic Diversity in the Butterfly Heliconius melpomene. Genetics, 203(1), 525–541. 10.1534/genetics.115.183285

Martini, S., Silvotti, L., Shirazi, A., Ryba, N. J. P., & Tirindelli, R. (2001). Co-Expression of Putative Pheromone Receptors in the Sensory Neurons of the Vomeronasal Organ. Journal of Neuroscience, 21(3), 843–848. 10.1523/JNEUROSCI.21-03-00843.2001

Meier, J. I., Marques, D. A., Wagner, C. E., Excoffier, L., & Seehausen, O. (2018). Genomics of Parallel Ecological Speciation in Lake Victoria Cichlids. Molecular Biology and Evolution, 35(6), 1489–1506. 10.1093/molbev/msy051

Meier, J. I., Sousa, V. C., Marques, D. A., Selz, O. M., Wagner, C. E., Excoffier, L., & Seehausen, O. (2017). Demographic modelling with whole-genome data reveals parallel origin of similar Pundamilia cichlid species after hybridization. Molecular Ecology, 26(1), 123–141. 10.1111/mec.13838

Nachman, M. W. (2002). Variation in recombination rate across the genome: Evidence and implications. Current Opinion in Genetics & Development, 12(6), 657–663. 10.1016/S0959-437X(02)00358-1

Nadeau, N. J., Whibley, A., Jones, R. T., Davey, J. W., Dasmahapatra, K. K., Baxter, S. W., Quail, M. A., Joron, M., ffrench-Constant, R. H., Blaxter, M. L., Mallet, J., & Jiggins, C. D. (2012). Genomic islands of divergence in hybridizing *Heliconius* butterflies identified by large-scale targeted sequencing. Philosophical Transactions of the Royal Society B: Biological Sciences, 367(1587), 343–353. 10.1098/rstb.2011.0198

Nei, M., & Rooney, A. P. (2005). Concerted and Birth-and-Death Evolution of Multigene Families. Annual Review of Genetics, 39(1), 121–152. 10.1146/annurev.genet.39.073003.112240

Niimura, Y., & Nei, M. (2007). Extensive Gains and Losses of Olfactory Receptor Genes in Mammalian Evolution. PLOS ONE, 2(8), e708. 10.1371/journal.pone.0000708

Noor, M. a. F., & Bennett, S. M. (2009). Islands of speciation or mirages in the desert? Examining the role of restricted recombination in maintaining species. Heredity, 103(6), 439–444. 10.1038/hdy.2009.151

Pamilo, P., & Nei, M. (1988). Relationships between gene trees and species trees. Molecular Biology and Evolution, 5(5), 568–583. 10.1093/oxfordjournals.molbev.a040517

Pedersen, B. S., & Quinlan, A. R. (2018). Mosdepth: Quick coverage calculation for genomes and exomes. Bioinformatics, 34(5), 867–868. 10.1093/bioinformatics/btx699

Pockrandt, C., Alzamel, M., Iliopoulos, C. S., & Reinert, K. (2020). GenMap: Ultra-fast computation of genome mappability. Bioinformatics, 36(12), 3687–3692. 10.1093/bioinformatics/btaa222

Quinlan, A. R., & Hall, I. M. (2010). BEDTools: A flexible suite of utilities for comparing genomic features. Bioinformatics, 26(6), 841–842. 10.1093/bioinformatics/btq033

Ravinet, M., Faria, R., Butlin, R. K., Galindo, J., Bierne, N., Rafajlović, M., Noor, M. A. F., Mehlig, B., & Westram, A. M. (2017). Interpreting the genomic landscape of speciation: A road map for finding barriers to gene flow. Journal of Evolutionary Biology, 30(8), 1450–1477. 10.1111/jeb.13047

Renaut, S., Grassa, C. J., Yeaman, S., Moyers, B. T., Lai, Z., Kane, N. C., Bowers, J. E., Burke, J. M., & Rieseberg, L. H. (2013). Genomic islands of divergence are not affected by geography of speciation in sunflowers. Nature Communications, 4(1), 1827. 10.1038/ncomms2833

Sanders, K. L., Lee, M. S. Y., Mumpuni, Bertozzi, T., & Rasmussen, A. R. (2013). Multilocus phylogeny and recent rapid radiation of the viviparous sea snakes (Elapidae: Hydrophiinae). Molecular Phylogenetics and Evolution, 66(3), 575–591. 10.1016/j.ympev.2012.09.021

Sanders, K. L., Mumpuni, & Lee, M. S. Y. (2010). Uncoupling ecological innovation and speciation in sea snakes (Elapidae, Hydrophiinae, Hydrophiini). Journal of Evolutionary Biology, 23(12), 2685–2693. 10.1111/j.1420-9101.2010.02131.x

Sanders, K. L., Rasmussen, A. R., Mumpuni, Elmberg, J., de Silva, A., Guinea, M. L., & Lee, M. S. Y. (2013). Recent rapid speciation and ecomorph divergence in Indo-Australian sea snakes. Molecular Ecology, 22(10), 2742–2759. 10.1111/mec.12291

Sayers, E. W., Bolton, E. E., Brister, J. R., Canese, K., Chan, J., Comeau, D. C., Connor, R., Funk, K., Kelly, C., Kim, S., Madej, T., Marchler-Bauer, A., Lanczycki, C., Lathrop, S., Lu, Z., Thibaud-Nissen, F., Murphy, T., Phan, L., Skripchenko, Y.,…Sherry, S. T. (2022). Database resources of the national center for biotechnology information. Nucleic Acids Research, 50(D1), D20–D26. 10.1093/nar/gkab1112

Schield, D. R., Carter, J. K., Alderman, M. G., Farleigh, K., Highland, D. K., & Safran, R. J. (2025). Recombination Rate and Recurrent Linked Selection Shape Correlated Genomic Landscapes Across a Continuum of Divergence in Swallows. Molecular Ecology, 34(22), e70074. 10.1111/mec.70074

Schiffels, S., & Wang, K. (2020). MSMC and MSMC2: The Multiple Sequentially Markovian Coalescent. In J. Y. Dutheil (Ed.), Statistical Population Genomics (pp. 147–166). Springer US. 10.1007/978-1-0716-0199-0_7

Schwenk, K. (1994). Why Snakes Have Forked Tongues. Science, 263(5153), 1573–1577. 10.1126/science.263.5153.1573

Seehausen, O., Butlin, R. K., Keller, I., Wagner, C. E., Boughman, J. W., Hohenlohe, P. A., Peichel, C. L., Saetre, G.-P., Bank, C., Brännström, Å., Brelsford, A., Clarkson, C. S., Eroukhmanoff, F., Feder, J. L., Fischer, M. C., Foote, A. D., Franchini, P., Jiggins, C. D., Jones, F. C.,…Widmer, A. (2014). Genomics and the origin of species. Nature Reviews Genetics, 15(3), 176–192. 10.1038/nrg3644

Shang, H., Field, D. L., Paun, O., Rendón-Anaya, M., Hess, J., Vogl, C., Liu, J., Ingvarsson, P. K., Lexer, C., & Leroy, T. (2023). Drivers of genomic landscapes of differentiation across a Populus divergence gradient. Molecular Ecology, 32(15), 4348–4361. 10.1111/mec.17034

Sherratt, E., Crowe-Riddell, J., Palci, A., Ammresh, Hutchinson, M. N., Lee, M. S. Y., & Sanders, K. L. (2025). Rapid evolution and cranial morphospace expansion during the terrestrial to marine transition in elapid snakes. Evolution, 79(12), 2629–2642. 10.1093/evolut/qpaf180

Sherratt, E., Rasmussen, A. R., & Sanders, K. L. (2018). Trophic specialization drives morphological evolution in sea snakes. Royal Society Open Science, 5(3), 172141. 10.1098/rsos.172141

Shine, R. (2005). All at sea: Aquatic life modifies mate-recognition modalities in sea snakes (Emydocephalus annulatus, Hydrophiidae). Behavioral Ecology and Sociobiology, 57(6), 591–598. 10.1007/s00265-004-0897-z

Shine, R., Shine, T., & Goiran, C. (2019). Morphology, reproduction and diet of the greater sea snake, Hydrophis major (Elapidae, Hydrophiinae). Coral Reefs, 38(5), 1057–1064. 10.1007/s00338-019-01833-5

Silva, L., & Antunes, A. (2017). Vomeronasal Receptors in Vertebrates and the Evolution of Pheromone Detection. Annual Review of Animal Biosciences, 5(Volume 5, 2017), 353–370. 10.1146/annurev-animal-022516-022801

Smith, J. M., & Haigh, J. (1974). The hitch-hiking effect of a favourable gene. Genetics Research, 23(1), 23–35. 10.1017/S0016672300014634

Steinegger, M., & Söding, J. (2017). MMseqs2 enables sensitive protein sequence searching for the analysis of massive data sets. Nature Biotechnology, 35(11), 1026–1028. 10.1038/nbt.3988

Tarasov, A., Vilella, A. J., Cuppen, E., Nijman, I. J., & Prins, P. (2015). Sambamba: Fast processing of NGS alignment formats. Bioinformatics, 31(12), 2032–2034. 10.1093/bioinformatics/btv098

The UniProt Consortium. (2025). UniProt: The Universal Protein Knowledgebase in 2025. Nucleic Acids Research, 53(D1), D609–D617. 10.1093/nar/gkae1010

Thorvaldsdóttir, H., Robinson, J. T., & Mesirov, J. P. (2013). Integrative Genomics Viewer (IGV): High-performance genomics data visualization and exploration. Briefings in Bioinformatics, 14(2), 178–192. 10.1093/bib/bbs017

Turner, T. L., Hahn, M. W., & Nuzhdin, S. V. (2005). Genomic Islands of Speciation in Anopheles gambiae. PLOS Biology, 3(9), e285. 10.1371/journal.pbio.0030285

Ukuwela, K. D. B., de Silva, A., Mumpuni, Fry, B. G., & Sanders, K. L. (2014). Multilocus phylogeography of the sea snake Hydrophis curtus reveals historical vicariance and cryptic lineage diversity. Zoologica Scripta, 43(5), 472–484. 10.1111/zsc.12070

Vasimuddin, Md., Misra, S., Li, H., & Aluru, S. (2019). Efficient Architecture-Aware Acceleration of BWA-MEM for Multicore Systems. 2019 IEEE International Parallel and Distributed Processing Symposium (IPDPS), 314–324. 10.1109/IPDPS.2019.00041

Voris, H. K. (2000). Maps of Pleistocene sea levels in Southeast Asia: Shorelines, river systems and time durations. Journal of Biogeography, 27(5), 1153–1167. 10.1046/j.1365-2699.2000.00489.x

Voris, H. K., & Voris, H. H. (1983). Feeding Strategies in Marine Snakes: An Analysis of Evolutionary, Morphological, Behavioral and Ecological Relationships. American Zoologist, 23(2), 411–425. 10.1093/icb/23.2.411

Wang, J., & Whitlock, M. C. (2003). Estimating Effective Population Size and Migration Rates From Genetic Samples Over Space and Time. Genetics, 163(1), 429–446. 10.1093/genetics/163.1.429

Watterson, G. A. (1975). On the number of segregating sites in genetical models without recombination. Theoretical Population Biology, 7(2), 256–276. 10.1016/0040-5809(75)90020-9

Wolf, J. B. W., & Ellegren, H. (2017). Making sense of genomic islands of differentiation in light of speciation. Nature Reviews Genetics, 18(2), 87–100. 10.1038/nrg.2016.133

Wood, D. E., Lu, J., & Langmead, B. (2019). Improved metagenomic analysis with Kraken 2. Genome Biology, 20(1), 257. 10.1186/s13059-019-1891-0

Wu, C.-I., & Ting, C.-T. (2004). Genes and speciation. Nature Reviews Genetics, 5(2), 114–122. 10.1038/nrg1269

Wynn, E. H., Sánchez-Andrade, G., Carss, K. J., & Logan, D. W. (2012). Genomic variation in the vomeronasal receptor gene repertoires of inbred mice. BMC Genomics, 13(1), 415. 10.1186/1471-2164-13-415

Zhang, C., Dong, S.-S., Xu, J.-Y., He, W.-M., & Yang, T.-L. (2019). PopLDdecay: A fast and effective tool for linkage disequilibrium decay analysis based on variant call format files. Bioinformatics, 35(10), 1786–1788. 10.1093/bioinformatics/bty875

Zhu, Z., Dobry, J., Wapstra, E., Zhou, Q., & Ezaz, T. (2026). Gene traffic mediated by transposable elements shaped the dynamic evolution of ancient sex chromosomes of varanid lizard. Journal of Genetics and Genomics, 53(3), 497–509. 10.1016/j.jgg.2025.08.002

